# Post-transcriptional regulator HuR promotes immune evasion in pancreatic ductal adenocarcinoma

**DOI:** 10.1101/2025.02.07.632847

**Authors:** Yifei Guo, Jennifer M. Finan, Alexandra Q. Bartlett, Shamilene Sivagnanam, Katie E. Blise, Nell Kirchberger, Konjit Betre, Grace A. McCarthy, Kevin Hawthorne, Canping Chen, Aaron Grossberg, Zheng Xia, Lisa M. Coussens, Rosalie C. Sears, Jonathan R. Brody, Robert Eil

## Abstract

The tumor microenvironment (TME) of pancreatic ductal adenocarcinoma (PDAC) is characterized by a limited infiltration of tumor-specific T cells and anti-tumor T cell activity. Extracellular factors in the PDAC TME have been widely reported to mediate immune suppression, but the contribution from tumor-intrinsic factors is not well understood. The RNA-binding protein, HuR (*ELAVL1*), is enriched in PDAC and negatively correlates with T cell infiltration. In an immunocompetent Kras-p53-Cre (KPC) orthotopic model of PDAC, we found that genetic disruption of HuR impaired tumor growth due to a novel role of HuR inducing T-cell suppression. Importantly, we found that HuR depletion in tumors enhanced both T cell number and activation states and diminished myeloid phenotypes by comprehensive spatial profiling of the PDAC TME. Mechanistically, HuR mediated the stabilization of mTOR pathway transcripts, and inhibition of mTOR activity rescued the impaired function of local T cells. Translating these findings, we demonstrated that HuR depletion sensitized PDAC tumors to immune checkpoint blockade, while isogenic, wildtype tumors are resistant. For the first time, we show that HuR facilitates tumor immune suppression in PDAC by inhibiting T cell infiltration and function and implicate targeting HuR as a potential therapeutic strategy in combination with immunotherapy.

**SIGNIFICANCE:** This study identified a novel mechanism that HuR supports pancreatic tumor growth by restricting T cell infiltration, promoting immune evasion. Our work supports targeting strategies against HuR in PDAC with the goal of enhancing PDAC sensitivity to immune-based cancer therapies, such as checkpoint blockade and T cell transfer.

## INTRODUCTION

Pancreatic ductal adenocarcinoma (PDAC) is one of the leading causes of cancer-related death worldwide with a 5-year survival rate of 13%, underscoring the need to develop improved therapeutic strategies^1^. The complex tumor microenvironment (TME) of PDAC is thought to hinder both chemo- and immunotherapy efficacy^2^. Despite the presence of immunogenic neoantigens that are predicted to be recognized by endogenous T cells, PDAC has remained resistant to immune checkpoint blockade (ICB)^3–5^. Some have proposed that the ability of cancer cells to downregulate antigen presentation machinery is a major driver of evading T cell recognition in PDAC. However, only a small proportion (∼1-2%) of PDAC contain mutations within antigen presentation or processing machinery^6^. Even in patients with mismatch repair deficient (MMR-D) driven PDAC, where neoantigen frequency is equivalent to other MMR-D cancers, ICB had compromised rates of objective response in PDAC in comparison to other MMR-D cancers^7^. Thus, we and others have concluded that unresolved tumor-intrinsic sources of T cell suppression exist in PDAC. Specifically, cytotoxic T cells in the PDAC TME have characteristics of T cell dysfunction, such as an inadequate response to stimulation (hypo-responsiveness), which is recognized as a major source of tumor immune escape^8–10^. Therefore, in PDAC, there remains a need to understand and reverse the tumor specific mechanisms that drive resistance to cancer immunotherapies^11^.

Investigators continue to appreciate that TME features, including cytokines and cell-cell interactions, can restrain the antitumor functions of CD4^+^ and CD8^+^ T cells. In tumors, PDAC especially, a combination of rapid cellular proliferation and pathologic extracellular architecture contribute to a microenvironment that is low in both glucose and oxygen^12,13^. Meanwhile, T cells depend on consumption of nutrients to effectively respond to TCR ligation with acquisition of effector programs, production of cytokines, and target cell cytolysis. Thus, the tumor-imposed nutrient deficit results in a metabolic competition with local immune cells, contributing to T cell hypo-responsiveness and resistance to immunotherapies^14^. As others have increasingly appreciated the importance of nutrient availability and metabolism for effective T cell antitumor function, interest continues to grow in therapies that could increase nutrient availability and consumption by tumor-specific T cells^15^.

In human PDAC and preclinical models, the RNA-binding protein, human antigen R (HuR, *ELAVL1*) is known to be hyperactive, supporting multiple programmatic hallmarks of cancer^12,13,16,17^. Functionally, HuR recognizes and binds to 3’ untranslated regions (UTRs) of transcripts, enhancing their stability and translation^18^. For instance, HuR reshapes the metabolic profile of PDAC, in part, by stabilizing *IDH1* transcripts and reprogramming both glycolytic and lipogenic metabolism^13,19,20^. Broadly, HuR activity is associated with Myc programs and glycolytic metabolism, enhancing the activation of c-Myc programs and consumption of surrounding nutrients for growth in size and cell division^12,21,22^. Additionally, pancreas-specific HuR overexpression alone caused fibro-inflammation in the mouse pancreas and led to an increase in precursor lesions of PDAC in cooperation with a *Kras^G12D^*mutation^23^. Our prior observations support a model wherein tumor-intrinsic HuR can mediate cell-cell communication and alter the PDAC TME in immune compromised models^13,24^. In the context of metabolic competition between tumor and immune cells, we hypothesized that tumor-intrinsic HuR activity may drive a competitive metabolic advantage over neighboring T cells. Yet, due to a lack of murine PDAC model that can capture the immune contexture with HuR depletion, the role of HuR in the PDAC-immune crosstalk has not been examined^16,17,24^.

Taken together, our prior works point to a hypothesis that HuR may have a substantial role in mediating immune evasion in the PDAC TME. Moreover, we report here that HuR expression is negatively correlated with both T cell infiltration and outcome in PDAC patients. Thus, we investigated whether depletion of HuR in PDAC would impact tumor growth and anti-tumor immune activity in an immunocompetent model of PDAC. To this end, we developed an isogenic *Kras^G12D^,Trp53^R172H^,Pdx1-Cre* (KPC) mutant-driven murine PDAC model, with and without HuR, to further elucidate the impact of tumor-intrinsic HuR on the immune components of the PDAC TME^25^. We observed that tumors with disruption in the *Elavl1* locus (HuR) had impaired tumor growth *in vivo.* We then assayed the tumors with multiplex immunohistochemistry (mIHC)^26^, a single-cell spatial proteomics imaging technology, and subsequently used machine learning to unbiasedly detect the main differences in TME immune contexture between tumors with HuR versus tumors without HuR. We found that HuR-deficient KPC tumors were most distinctly characterized by their increased infiltration of CD4^+^ T cells and increased frequency of spatial proximities involving anti-tumoral “M1-like” CD206^-^ macrophages and CD4^+^ T cells, while HuR-competent KPC tumors possessed an immunosuppressive, myeloid-enhanced spatial phenotype. Moreover, antibody-based depletion of CD4^+^ and CD8^+^ T cells restored the *in vivo* growth of HuR-deficient KPC tumors, demonstrating that HuR is required for intact PDAC growth in the immunocompetent setting. Introduction of a model antigen and assessment of tumor specific T cells identified that tumor-intrinsic HuR deterred the infiltration of antigen-specific CD8^+^ T cells and their acquisition of effector functionality. Mechanistically, HuR mediated stabilization of mTOR pathway-related transcripts and enhanced PDAC nutrient consumption to the detriment of local T-cell antitumor function. Consistent with such a model, inhibition of the mTOR pathway in PDAC cells enhanced the cytokine production capacity of co-cultured CD8^+^ T cells in an HuR-dependent manner. Therapeutically, KPC tumor’s resistance to ICB treatment was dependent on the presence of HuR, and HuR depletion combined with ICB led to tumor regression. Herein, we provide a deeper understanding of a potent immune suppression mechanism driven by the pro-oncogenic protein, HuR, in PDAC.

## METHODS

### HuR signature scoring of RNA-seq OPTR cohort and survival analysis

Expression of the HuR activity signature^17^ was assessed for 218 primary PDAC tumor RNA-seq profiles from the Oregon Pancreas Tissue Registry (OPTR) cohort described in Link JM et al.^27^. Pseudoaligned transcript abundances for each RNA-seq profile were summed to gene level using tximport^28^ and subsequently used as input for DESeq2^29^. Gene expression values were transformed with DESeq2 variance-stabilizing transformation (VST). The GSVA^30^ algorithm was then applied to VST count data to assign HuR signature scores to each sample. Overall survival was assessed in the same 208 patients described previously^27^ but filtered by HuR signature scores. Patients were stratified into groups representing high HuR signature expression (GSVA score in the top quartile) and low (GSVA score in the bottom quartile). Survival was compared using the survival^31^ and survminer^32^ packages.

### Single-cell data collection and analysis

Single-cell RNA sequencing (scRNA-seq) data were sourced from the Deeply Integrated Single-Cell Omics (DISCO) database^33^. A total of 115 PDAC samples, encompassing 557,304 cells, were utilized to investigate the relationship between *ELAVL1* gene expression in malignant cells and lymphocyte infiltration. Malignant cells were identified using the R package scATOMIC (v2.0.2)^34^ with the parameter “known_cancer_type = PAAD cell” and default settings for other parameters. Spearman’s correlation analysis was conducted to assess the association between gene expression levels in tumor cells and cell-type infiltration in each sample.

### Immunohistochemistry staining of tumors

Mouse tissues were fixed in 4% PFA overnight at 4°C before embedding and 5 µm sectioning. As previously described^24^, paraffin-embedded histological sections were stained for HuR (Santa Cruz Biotechnology; sc-5261) overnight at 4°C, followed by incubation with a biotinylated secondary antibody for 1 hour at room temperature. Color was developed with DAB substrate (Vector Laboratories; SK-4100), and mount for imaging. Brightfield images were acquired by Zeiss Axio slide scanner.

### Cell lines and virus

The PDAC murine cell line, KPC8069 (Kras^G12D^-Trp53^R172H^-Pdx1Cre), was a gift from Dr. Michael Hollingsworth. Cells were cultured in normal DMEM with 10% fetal bovine serum (Gibco) and 1% penicillin/streptomycin (Thermo Scientific; 15070063). Cell lines were maintained at 37°C in 5% humidified CO_2_. Cell lines were *Mycoplasma* tested routinely using Mycoplasma Detection Kit (SouthernBiotech; 13100-01). Before any experiment, cells were passaged at least twice after thawing.

For HuR re-expression, KPC8069 cells were transfected with ecotropic retrovirus either carrying codon-enhanced *Elavl1* sequence or empty vector and supplemented with 1 μg/ml polybrene (Millipore Sigma; TR1003G). Cells containing the target vectors were selected by hygromycin (Millipore Sigma; H3274) and validated for HuR expression with immunoblotting. For ovalbumin (OVA) expression, KPC8069 cells were transfected with ecotropic retrovirus carrying the full-length OVA sequence and supplemented with 1 μg/ml polybrene. Cells containing the target vectors were selected by blasticidin (Alfa Aesar; J67216-XF) and validated for OVA expression with flow cytometry.

All ecotropic retroviruses are generated by packaging 20 μg of plasmid DNA synthesized by Genscript, Inc. containing genes of interest in an MSCV derived γ-retrovirus backbone (derived from Addgene; 122727) along with 3 μg pCL-Eco plasmid DNA (Addgene; 12371) with platinum-E ecotropic packaging cells (Cell Biolabs; RV-101) and jetOPTIMUS transfection reagent (Avantor; 101000006).

### INDEL assessment

For INDEL assessment, genomic DNA was harvested from cells at time of analysis using the DNeasy Blood and Tissue DNA kit (Qiagen; 69504), and the region containing *Elavl1* short guide (sg) RNA target was PCR-amplified and Sanger sequenced by Eurofins Genomics. For each sample, a 493-bp region was amplified for sequencing (forward primer: 5′-ACTGGGGCTGAAGAAAAGTAGT-3′, reverse: 5′-GAACTGAATCATCTCCCCGGT-3′). 100 to 200 ng of each amplicon was sent for sequencing with 2 pmol of its respective sequencing primer (5′-TAGTCTGAGGTTAAAGAACAAAACTTCATT-3′). The indel percentage of amplicons derived from HuR-KO cells was deduced using the Synthego ICE tool (Synthego; ICE Analysis v3.0).

### Immunoblot analysis

Cells were lysed using ice-cold RIPA buffer (Thermo Scientific; 89900) supplemented with protease inhibitors (Thermo Scientific; 87786). Protein concentration was quantified using the Pierce BCA Protein Assay kit (Thermo Scientific; 23225). Equal amounts of whole protein extracts were loaded and separated by electrophoresis on a 4% to 12% Bis-Tris gel (Invitrogen; NP0321BOX) and transferred to a PVDF membrane (Bio-Rad; 1704156). Blots were blocked in Intercept® (TBS) Blocking Buffer (LI-COR Biosciences; 927-70003) and then probed with antibodies against anti-HuR (1:1000, Santa Cruz Biotechnology; sc-5261) and anti-α-tubulin (1:5000, Invitrogen; A11126), followed by Thermo Fisher Scientific Alexa secondary antibodies. Immunofluorescence was measured using a digital imager (Invitrogen; iBright 1500).

### RNA extraction and RT-qPCR

RNA was extracted using RNeasy Mini Kit (Qiagen; 74106). RNA concentrations and quality were assessed by NanoDrop spectrophotometer (Thermo Scientific). Complementary DNA (cDNA) was made from 500 ng RNA using High Capacity cDNA Reverse Transcriptase Kit (Thermo Scientific; 4368813). RT-qPCR was performed with SYBR green master mix (Thermo Scientific; 4385617) with HuR primers and cDNA. Amplification signal was captured by RT-qPCR machine (Thermo Scientific; QuantStudio 7 Flex). Relative quantifications were assessed using the Log_2_–ΔΔCt method, using GAPDH for normalization.

### Mouse experiments

Animal protocol # IP00003322 and #IP00003155 were approved by Institutional Animal Care Regulations and Use Committee of OHSU. These protocols are specifically applicable to the experiments reported in this manuscript. All mice were obtained from The Jackson Laboratory. C57BL/6 mice (Cat. 000664) of 8–9 weeks of age were used as recipient hosts for KPC tumor implantation. OT-I transgenic mice were used as OT-I T cell donors.

For T cell depletion studies, mice were intraperitoneally injected with αCD8 blocking antibody (200 μg, BioXcell; BE0061) and αCD4 blocking antibody (200 μg; BioXcell; BE0003-1) two days before tumor implantation. The treatment was maintained for the duration of the experiment by injecting every four days of αCD8 and αCD4 (100 μg each). For isotype controls, rat IgG2b (BioXcell; BE0090) at matching amount was used. This approach achieved >99% depletion of CD8^+^ and CD4^+^ T cells in spleens of T cell-depleted mice compared to control mice at endpoint.

For αPD-1 treatment, mice were intraperitoneally injected with αPD-1 blocking antibody (200 μg, BioXcell; BP0146) or isotype controls, rat IgG2a (BioXcell; BE0089) every 4 days. The first injections happened 5 days after tumor implantation, and each mouse received a total of 4 injections.

### Mouse PDAC tumor model

For implantable tumor experiments, KPC tumor cells (1 × 10^6^) were resuspended in 100 μl of 1:1 mixture of PBS (Fisher Scientific; 14-190-250) and Matrigel (Fisher Scientific; 354234), injected subcutaneously into the flanks of mice, and allowed to grow for 37 days. Tumor length and width were measured by caliper every four days.

For orthotopic tumor growth experiments, KPC tumor cells (4 × 10^4^) were resuspended in 20 μl of 1:1 mixture of PBS and Matrigel, orthotopically injected into the tail of mouse pancreas, and allowed to grow for 20 days. For orthotopic tumor αPD-1 experiments, KPC tumor cells (9 × 10^4^) were injected and allowed to grow for 15 days. Earlier endpoint criteria included severe weight loss, or extreme weakness and inactivity. Tumor weight was measured by chemical scale (Sartorius) at the endpoint.

### Flow cytometry and analysis

Following euthanasia, tumors and spleens were promptly dissected, weighed, and stored in ice-cold PBS. Tumors were subsequently minced with surgical scissors and dissociated with GentleMACS octo-dissociator and preset program (m_impTumor_01, Miltenyi Biotec). The dissociated tumor was then filtered through a 40 μm filter. Spleens were mashed and filtered through a 40 μm filter. All samples were ACK lysed for 60 seconds using ACK lysing buffer (Gibco; A1049201), and then consolidated for staining.

Single-cell suspension of each sample was incubated with Live/Dead Fixable Aqua (Thermo Scientific; L34966) and Fc block (BD; 553142) in PBS for 30 min at 4°C, followed by incubation with primary fluorochrome-labeled antibodies at 4°C for 30 min in PBS with 0.5% BSA and 2 mM EDTA. For *ex vivo* cytokine production by T cell, samples were incubated for 4 hours at 37°C with PMA (20 ng/mL, Sigma; P1585), ionomycin (1 μg/mL, Sigma; I0634), and GolgiStop (BD; 554724). Intracellular staining was done using the Fixation/Permeabilization Kit (BD; 554722). Antibodies used in flow analysis are listed in Supplementary Table 1. Flow cytometric analysis was performed on Cytek Aurora spectral cytometer. Collected data were analyzed using FlowJo v10 (TreeStar).

### mIHC image acquisition and analysis

As previously described^10,35,36^, multiplex staining was performed on 5 μm sections and each stained image was scanned at 20× magnification on an Aperio AT2 scanner (Leica Biosystems). The antibody panel used consisted of lineage markers and phenotype markers (Supplementary Table 2). Multiple 2.0 mm^2^ tissue regions were sampled from each tumor, resulting in 31 total tissue regions across 8 tumors. Tissue regions were registered using Matlab Computer Vision Toolbox (The Mathworks, Inc.), color deconvolution was performed using ImageJ, and nuclei segmentation was performed using StarDist. Single-cell mean intensity for each stain was quantified using Cell Profiler^37^. Single biomarker positivity thresholds were set using FCS Express Image Cytometry RUO (De Novo Software). Single-cell classification was performed using R Statistical Software according to the defined phenotypic gating hierarchy (Supplementary Table 3).

### mIHC spatial quantifications

The spatial organization of cells within tissue regions was quantified via cell-cell spatial proximities and recurrent cellular neighborhoods (RCNs), as previously published^10^. Cell-cell spatial proximities were calculated by counting the frequency at which two cell types occur within 20 µm of each other, then dividing the count by the summed densities of the two cell types involved in the proximity per tissue region. This normalization was used to mitigate the effect of cells present in high densities. RCNs were calculated by first identifying neighborhoods for every cell present across all tissue regions. Neighborhoods were defined by counting all cells within a 60 µm radius from the given seed cell. 76 cells had no neighboring cells within 60 µm and were dropped from the RCN analysis. The remaining 429,602 cells across the dataset were grouped using k-means clustering according to the normalized phenotypic composition of their neighborhood. The elbow method was used to determine the optimal number of groups to cluster the cell neighborhoods into. This resulted in each cell being assigned to one of eight RCNs, and the proportions of RCNs present per tissue region were calculated. Previously published code^10^ was used to complete the spatial analyses using Python software and the following packages: pandas, numpy, scipy, and scikit-learn.

### Machine learning pipeline

Following the previously published pipeline^10^, an elastic net model was trained to classify samples based on tumor genotype (*scramble* vs *sgElavl1*). Input features to the model included 12 cell population densities, 78 cell-cell proximity frequencies, and 8 RCN proportions, resulting in 98 total TME features calculated for each of the 31 tissue regions. Features were log10+1 transformed and scaled from zero to one to improve model interpretability. Predictions were made on a tissue region basis to maximize sample size and provide the model with the greatest amount of data to learn biologically relevant patterns. A leave-one-tumor-out cross-validation approach was used to prevent data leakage between cross-validation folds, and model hyperparameters remained the same across folds to prevent artificial inflation of performance. Briefly, for each fold, a new model was trained from tissue regions from seven of the eight tumors, while tissue regions from the remaining one tumor were withheld and used to test the model. Within each fold, the train set was balanced using Synthetic Minority Oversampling Technique (SMOTE) to upsample the minority class to equal the majority class^38^. This process was repeated until tissue regions from all 8 tumors appeared in the test set. Test set predictions were aggregated across the 8 folds into one confusion matrix, and final performance metrics were calculated from this confusion matrix. Coefficients were recorded for each of the 8 models and then averaged across all models to calculate final feature importance. Previously published code^10^ was used to perform the machine learning pipeline using Python software and the following packages: pandas, numpy, scikit-learn, and imbalanced-learn. The elastic net classifier was built using scikit-learn’s LogisticRegression function with the ‘penalty’ and ‘l1_ratio’ parameters set to ‘elasticnet’ and 0.5, respectively.

### In vitro T cell activation and co-cultures

Murine OT-I CD8^+^ T cells were isolated from the spleens of OT-I transgenic mice (The Jackson Laboratory; #003831) using the EasySep Mouse CD8^+^ T cell Isolation Kit (Stemcell; 19853), which yielded an enrichment of 96%– 100% purity. Isolated CD8^+^ T cells were stimulated *in vitro* with plate-bound anti-CD3 and anti-CD28 (5 μg/mL; BioXcell) for 24 hours and expanded in RPMI-1640 (Gibco) completed with 10% FBS (Gibco), 1% nonessential amino acids (Gibco), 1% sodium pyruvate (Gibco), 1% penicillin/streptomycin (Gibco), 57 μmol/L 2-BME (Sigma), and 100 IU/mL of recombinant human IL-2 (Peprotech).

For CD8^+^ T cell co-culture with tumor cells, 50,000 CD8^+^ T cells were incubated with 50,000 tumor cells in 96-well round-bottom plates for 24 hours at 37°C with phorbol 12-myristate 13-acetate (PMA) (20 ng/mL, Sigma; P1585), ionomycin (1 μg/mL, Sigma; I0634), and GolgiStop (BD; 554724). Intracellular staining was done using the Fixation/Permeabilization Kit and antibodies used are listed in Supplementary Table 1. SIINFEKL tetramers were obtained from the NIH tetramer core facility.

### RNA-sequencing

RNA was isolated from frozen cells using RNeasy Mini Kit (Qiagen; 74106). RNA concentrations and quality were assessed by NanoDrop spectrophotometer. Preparation of RNA library and transcriptome sequencing were conducted by Novogene Co., Ltd using the HiSeq Illumina platform. Novogene analysis service aligned raw reads to GRCm38 genome and performed DESeq2 differential expression analysis. Gene set enrichment analysis (GSEA) was done using GSEA v4.2.3 software with the MSigDB hallmark and curated gene sets^39,40^.

### Ribonucleoprotein immunoprecipitation assay

KPC cells were washed and harvested with cell scrapper in cold-PBS. Ribonucleoprotein immunoprecipitation assay was done according to manufacturer’s instructions (MilliporeSigma; 17-701) with modification to RNA purification. Lysates were immunoprecipitated by using 10 μg of either anti-HuR antibody (Santa Cruz Biotechnology; sc-5261) or anti-IgG control (MilliporeSigma; 12-371). For RNA purification from immunoprecipitation, the protocol was followed up to the precipitation of RNA in ethanol, after which the entire mixture was transferred to an RNA column using the RNeasy Micro Kit (Qiagen; 74004), and the manufacturer’s instructions were followed thereafter. RNA quality was analyzed via Bioanalyzer Pico chip by the Massively Parallel Sequencing Shared Resource Core at OHSU.

Preparation of RNA library and transcriptome sequencing were conducted by Novogene Co., Ltd using the HiSeq Illumina platform. Novogene analysis service aligned raw reads to GRCm38 genome and performed HuR-enriched peak annotation with MACS2 software^41^. KEGG (Kyoto Encyclopedia of Genes and Genomes) pathway enrichment analysis was done on HuR-enriched peaks.

### Statistical analysis

Data were analyzed using unpaired two-tailed Student’s *t*-test and one-way ANOVA test. In all cases, two-tailed tests with P values less than 0.05 were considered significant. Statistics were calculated using GraphPad Prism 10 software (GraphPad Software Inc). Results were graphed as mean ± standard deviation for *in vitro* experiments and ± standard error of mean for animal experiments. Experimental sample sizes were chosen using power calculations and preliminary experiments. Samples that had undergone technical failure during processing were excluded from subsequent analysis.

### Data availability

RNA-sequencing dataset is deposited at the GEO (Gene Expression Omnibus) database as GSE287307. RIP-sequencing dataset is deposited as GSE287349.

## RESULTS

### Tumor ELAVL1 (HuR) expression correlates with patient outcome and T cell infiltration in PDAC

We previously demonstrated that the RNA-binding protein HuR is highly expressed and activated (i.e. translocated to the cytoplasm) in PDAC compared to adjacent normal pancreas^24,42^. First, we applied a validated HuR activity signature highlighting HuR-dependent transcripts^17^ to a large RNA-sequencing dataset including surgical-resected PDAC patients^27^, and showed that a lower HuR target activity signature predicted significantly prolonged patient survival (**Fig. 1A**). Separately, baseline immune infiltration has been demonstrated to be associated with overall prognosis and response to ICB in a variety of cancers^43^. To further understand the relationship between HuR activity in PDAC and antitumor immune activity, we assayed DISCO single-cell RNA sequencing datasets and found tumor-intrinsic HuR expression levels negatively correlated with total CD3^+^ lymphocyte infiltration, i.e., lower tumor-intrinsic *ELAVL1* mRNA levels were associated with more abundant CD3^+^ lymphocyte infiltration in patient samples (**Fig. 1B** and **C**). To determine the causative relationship between tumor-intrinsic HuR and T cell infiltration, and antitumor function in PDAC, we used a syngenetic implantable mouse model of PDAC (KPC)^25^. To first assess whether HuR activity is recapitulated in our KPC model as in human PDAC, HuR protein expression and localization was visualized with IHC staining. In concordance with patient data, the KPC tumors had higher expression of whole-cell HuR compared to the healthy mouse pancreas (**Fig. 1D** and **E**). These findings support previous work that tumor-intrinsic HuR is a poor prognostic marker in PDAC and pro-oncogenic^17,44^, and in addition, negatively correlates with lymphocyte infiltration in PDAC. HuR is similarly elevated in the KPC model as in human PDAC providing a strong rationale for the investigation of tumor-intrinsic HuR in this immunocompetent murine PDAC model. To our knowledge, this is the first genetic assessment of the role of HuR in PDAC tumor *in vivo* in an immune intact setting.

**Figure 1.**
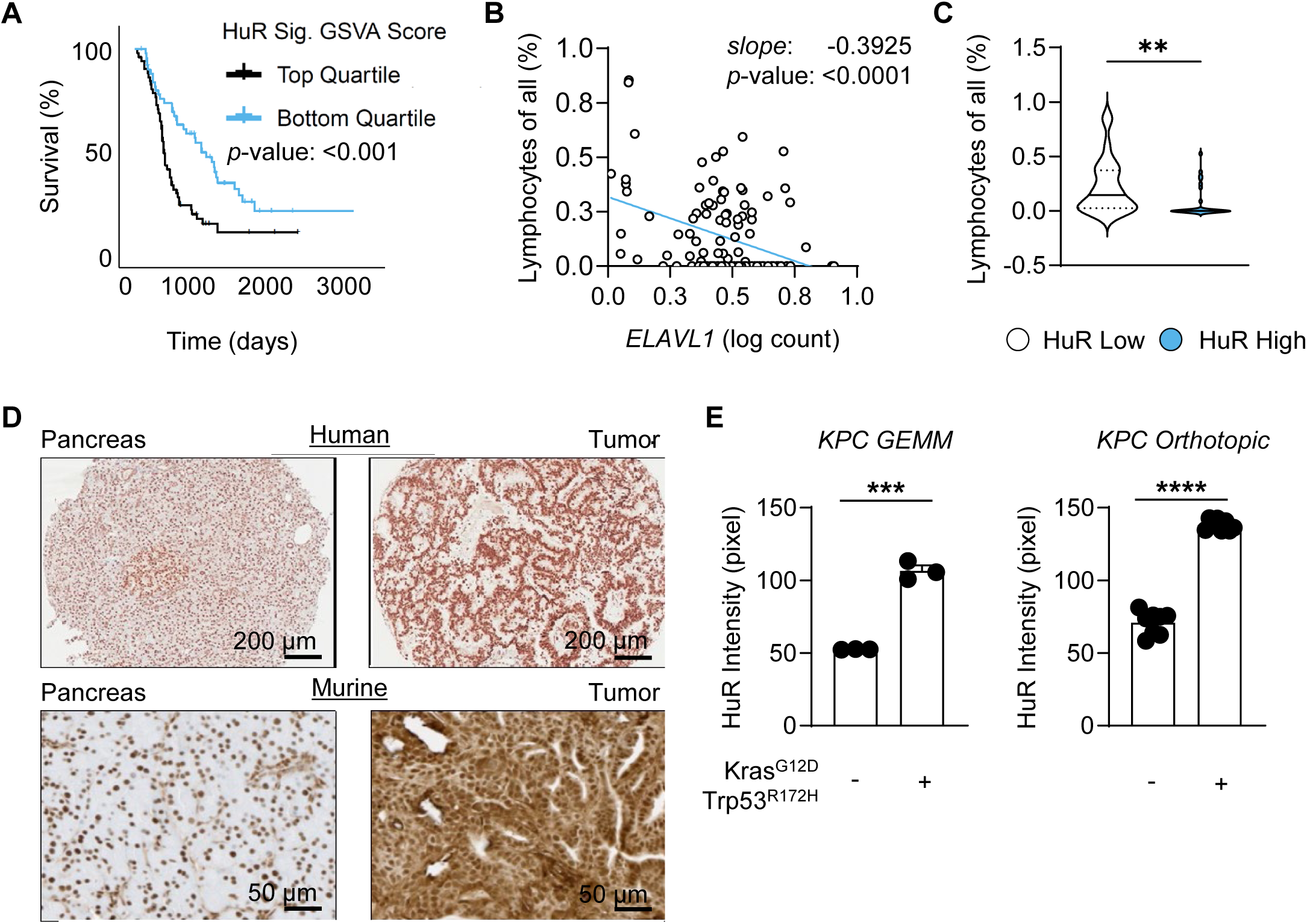
Tumor ELAVL1 (HuR) expression correlates with patient outcome and T cell infiltration in PDAC. (**A**) Kaplan-Meier plots demonstrating proportional overall survival in PDAC patients (n = 199) distinguished by low versus high HuR Activity Signature. (**B**) Scatter plot showing the Spearman’s correlation between ELAVL1 expression in tumor cells and lymphocyte infiltration, including CD8+ T cells, CD4+ T cells, and NK cells (n = 115 patients) and **(C)** violin plot demonstrating the difference in lymphocyte infiltration in HuR-low and HuR-high tumors from the same samples. (**D**) Representative images from patient tumor microarrays and KPC tumors stained for HuR expression and **(E)** quantification of HuR protein staining intensity of normal pancreas, KPC GEMM (n=3) and xenograft tumors (n = 8). Error bars represent standard error of the mean. ns, not significant; **P < 0.01; ***P < 0.005; ****P < 0.001, unpaired 2-tailed Student’s t tests **(C,E)**.

### Tumor-intrinsic HuR supports tumor growth in KPC model

We first established KPC cells with CRISPR-Cas9 mediated *Elavl1* genetic deletion using electroporation of scramble or *Elavl1* targeted sgRNAs complexed to purified recombinant Cas9 protein, as previously described^45^. We isolated and expanded single cell clones from these KPC HuR knockout (KO) pooled cell lines and three clones with stable knockout and proliferation performance were selected and used for subsequent analyses. We then validated and quantified CRISPR-Cas9 mediated disruption of the *Elavl1* locus via insertion or deletion (INDEL) via Sanger sequencing for our isolated clones (**Fig. 2A**). Accordingly, the clones had a significant reduction in HuR mRNA and lost HuR protein expression (**Fig. 2B** and **C**).

**Figure 2.**
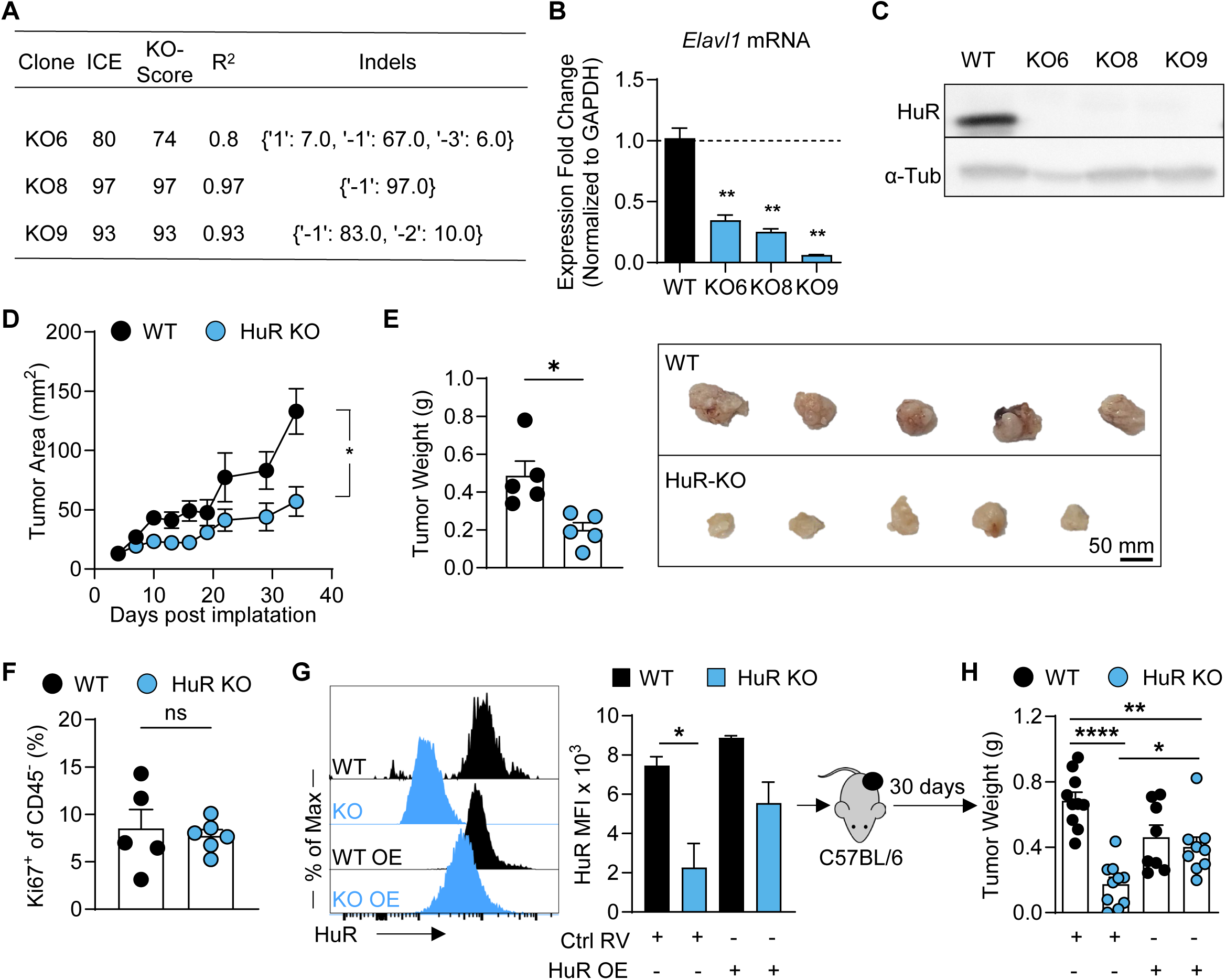
Tumor-intrinsic HuR supports tumor growth in KPC model. (**A**) INDEL quantification of single-cell clones isolated from CRISPR-Cas9 mediated disruption of the *Elavl1* locus in KPC8069 tumor cells and (**B**) quantification of mRNA abundance by RT-qPCR and (**C**) protein abundance by immunoblot of the same clones. (**D**) Rates of subcutaneous KPC tumor growth represented over time following implant of KPC WT and HuR-KO tumor cells (n = 5). (**E**) Summary quantification and pictures of the final tumor weight and size on day 34. (**F**) Quantification of the percentages of Ki67^+^ cells in total non-immune cells in subcutaneous KPC tumors by flow cytometry (n = 5). (**G**) Histogram and quantification of flow cytometry analysis of HuR expression in KPC WT and HuR-KO tumor cells retrovirally transduced with empty vector or HuR-overexpression vector prior to subcutaneous implantation to C57BL/6 hosts (n = 10). (**H**) Summary quantification of the final tumor weight on day 30. Error bars represent standard deviation (**B,F**) and standard error of the mean (**D,E,F,G,H**). *P < 0.05; **P < 0.01; ****P < 0.001, one-way ANOVA test (**B,G,H**), unpaired 2-tailed Student’s t tests (**D,E,F**).

To determine the impact of HuR on PDAC progression *in vivo*, we subcutaneously implanted KPC cells subjected to electroporation with scramble sgRNAs wildtype (WT) or sgRNAs targeting *Elavl1* (HuR-KO) into syngeneic C57BL/6 mice and tracked tumor growth over time. We found that the loss of tumor-intrinsic HuR significantly delayed growth over time by area and by end point weight (**Fig. 2D** and **E**). However, Ki67 staining showed no difference in tumor cell proliferation between these WT and HuR-KO tumors, suggesting that the decrease in tumor growth that resulted from the loss of PDAC HuR was dependent on a tumor-host interaction (**Fig. 2F**). Importantly, re-expression of the *Elavl1* sequence via durable retroviral transduction in these HuR-KO cells rescued the tumor growth defect (**Fig. 2F** and **G**). Cumulatively, these data causatively demonstrate, in genetic gain and loss of function, that HuR is required for PDAC progression *in vivo*.

### HuR-dependent growth of PDAC is immune mediated in vivo

Our compiled analysis of TCGA cohorts identified a negative correlation between *ELAVL1* mRNA abundance and total lymphocytes infiltration, suggesting that tumor-intrinsic HuR may act to constrain local immune activity and promote tumor progression. To define the functional relationship between endogenous antitumor immunity and tumor-intrinsic HuR, we orthotopically implanted KPC WT or HuR-KO into the pancreas of C57BL/6 hosts. We again observed a significant delay in growth in the HuR-KO tumors in comparison to KPC WT tumors in this physiological-relevant stromal and immune microenvironment (**Fig. 3A**). To provide a more granular understanding of the endogenous immune response, we performed enumerative and phenotypic assessment of intratumoral immune populations via flow cytometry. Our analysis revealed an increase in the abundance of CD4^+^ T cells in KPC HuR-KO tumors as compared to WT tumors (**Fig. 3B**). We also observed an increase in NK1.1^+^ T cells and CD19^+^ B cells as a proportion of all CD45^+^ leukocytes within HuR-KO tumors (Supplementary Fig. S1A). Phenotypic analysis of the tumor-infiltrating T cells demonstrated that both CD4^+^ and CD8^+^ T cells from HuR-KO tumors had a greater proportion expressing the transcription factor T-bet (*Tbx21*) (**Fig. 3C**). This is consistent with ongoing T cell effector function and polarization towards T helper Type 1 (T_H_1) effector specification, the cell population responsible for antitumor activity^46^. Additionally, the percentage of CD4^+^ Foxp3^+^ regulatory T (T_Reg_) cells trended lower in HuR-KO tumors compared to WT tumors, although not statistically significant (Supplementary Fig. S1B). Additionally, T cell surface activation phenotype markers CD25 and CD69 were more abundant on CD4^+^ and CD8^+^ T cells from HuR-KO tumors than WT tumors, while cell cycle marker Ki-67 stayed the same between groups (**Fig. 3D**). Surface expression of antigen presentation major histocompatibility complex (MHC) class I, H2-kb, and immune checkpoint protein, PD-L1, on tumor cells was not significantly different between WT and HuR-KO tumors, suggesting HuR mediates immune suppression in this model independently of antigen presentation or the expression of co-inhibitory surface ligands (Supplementary Fig. S1C). However, PD-L1 level was significantly higher in total HuR-KO cells than WT cells, indicating HuR-KO tumors were generally more immune activated (Supplementary Fig. S1D).

**Figure 3.**
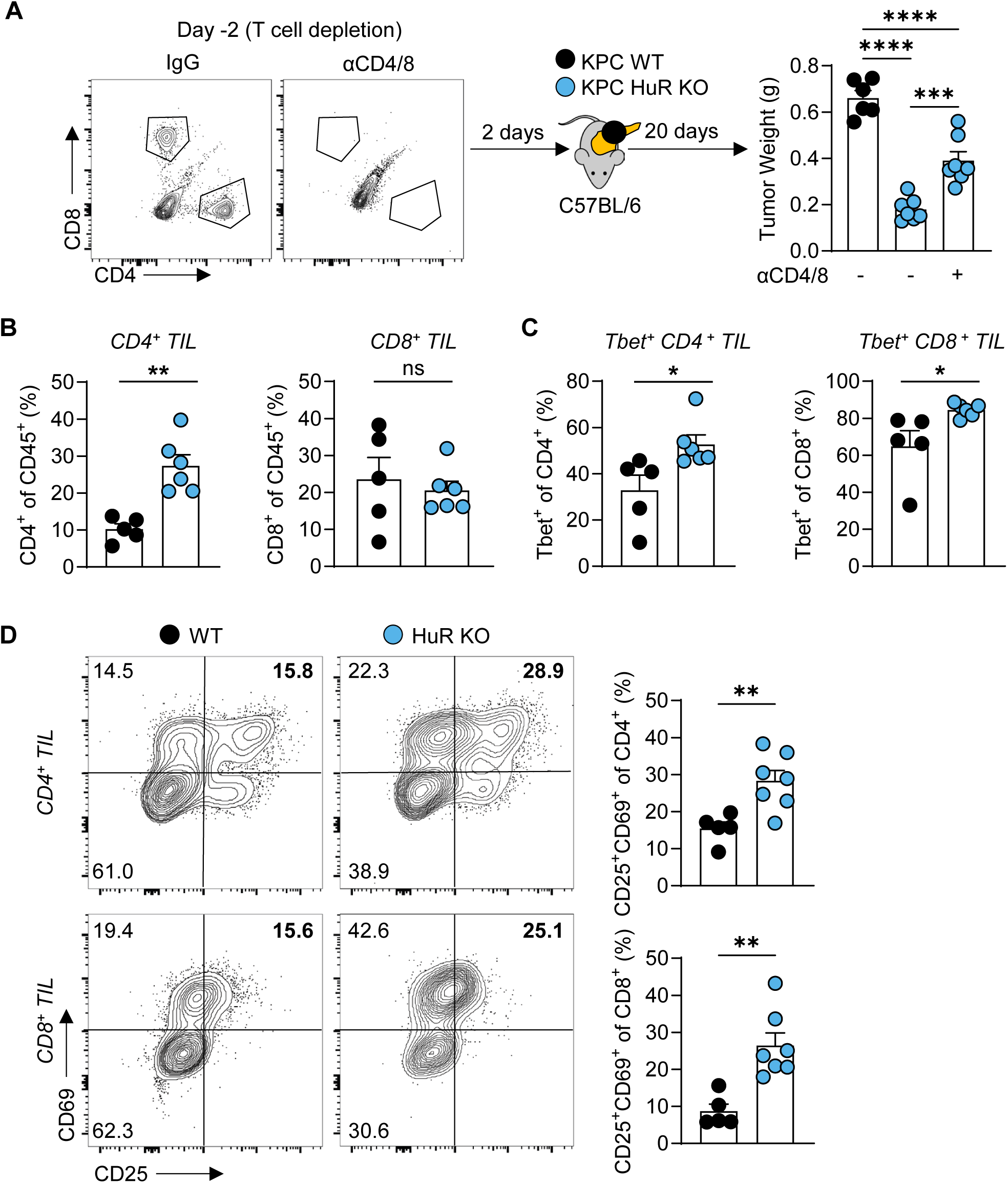
HuR-dependent growth of PDAC is immune mediated *in vivo*. (**A**) Representative flow cytometry illustrating enumerating T cells in host spleens treated with IgG or αCD4/8 before and during orthotopic tumor implantation and summary analysis of final KPC WT and HuR-KO tumor weight in host with or without T cell depletion (n = 6). (**B**) Summary analysis enumerating tumor infiltrating lymphocytes (TIL) in orthotopic KPC WT and HuR-KO tumors (n = 5). (**C**) Summary data depicting CD4 and CD8 T-cell Tbet^+^ percentages and (**D**) representative flow cytometry plot and summary data depicting CD25 and CD69 phenotypic states on CD4 and CD8 T cells as in (**B**). Error bars represent standard error of the mean. *P < 0.05; **P < 0.01; ***P < 0.005; ****P < 0.001, one-way ANOVA test (**A**), unpaired 2-tailed Student’s t tests (**B,C,D**).

Lastly, to test the causal relationship between endogenous antitumor immunity and HuR-mediated tumor progression, we performed antibody-mediated depletion of CD4^+^ and CD8^+^ T cells from C57BL/6 hosts followed by orthotopic implantation of KPC tumors with or without HuR depletion. We found that depletion of CD4^+^ and CD8^+^ T cells restored the tumor growth of HuR-KO tumors, suggesting that the defect in tumor growth of HuR-KO PDAC is due to a comparative increase in endogenous T cell-mediated tumor control in the absence of tumor-intrinsic HuR (**Fig. 3A**). Together these data indicate that the tumor regression in the KPC HuR-KO tumors is mediated by presence of activated T cells, which are otherwise suppressed in WT conditions.

### HuR limits T cell infiltration and activation, promoting an immunosuppressive and myeloid-driven TME landscape

Importantly, the TME is not just defined by the numerical presence of immune cells, but by how these cells are spatially organized, enabling communication with one another. Contrary to flow cytometry, mIHC preserves the spatial context of the tissue, providing deep phenotypic information and precise spatial locations of the cells present in the TME^26^. Prior research that utilized mIHC analysis has discovered novel cellular crosstalk and immunological niches that shape the progression of PDAC^10,35^. Therefore, we investigate how HuR impacts the PDAC TME spatial landscape to identify the key immunological crosstalk that led to HuR-mediated immune evasion. By performing mIHC on orthotopically implanted KPC WT and HuR-KO tumors, 429,669 cells were assayed in total across 31 2.0 mm^2^ tissue regions from 8 tumors (4 WT and 4 HuR-KO) by a 15-antibody mIHC panel (**Fig. 4A** and Supplementary Fig. S2A). Cells were hierarchically gated into 12 lineages, spanning lymphoid, myeloid, stromal, and neoplastic lineages (**Fig. 4B** and Supplemental Table 3). We then quantified TME cellular composition and spatial architecture in three ways, as previously described^10^: (i) cell density, (ii) cell–cell spatial proximities, and (iii) recurrent cellular neighborhoods (RCNs), resulting in 98 unique TME features calculated per tissue region. Because of the quantity and complexity of the mIHC data, we leveraged the previously published machine learning pipeline^10^ to first train an elastic net model to classify tumor genotype (WT vs HuR-KO) of a given tissue region based upon the 98 TME features, and then interrogate model coefficients as a means to perform feature selection and unbiasedly identify the top underlying TME immunobiology distinguishing KPC WT tumors from HuR-KO tumors.

**Figure 4.**
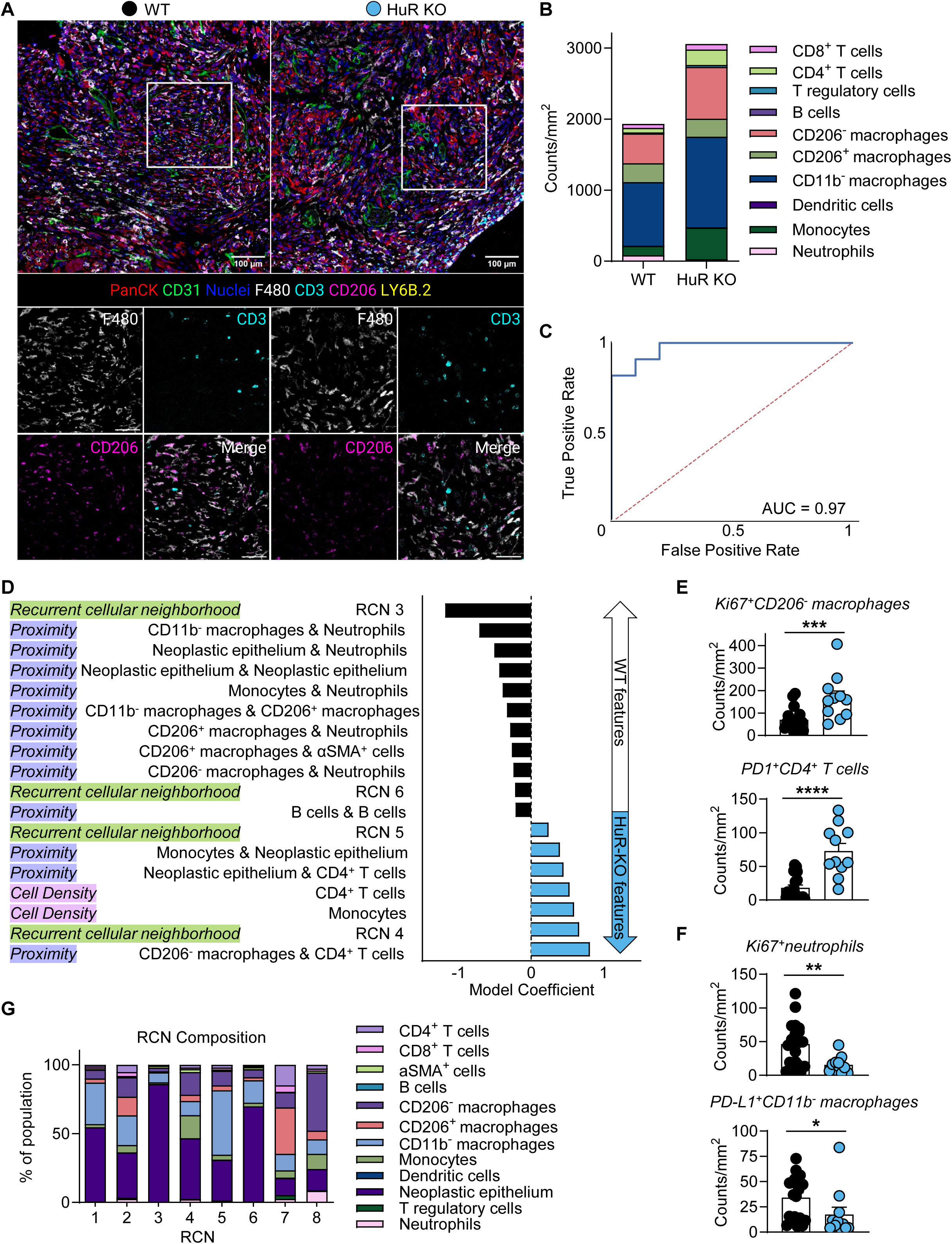
HuR limits T cell infiltration and activation, promoting an immunosuppressive and myeloid-driven TME landscape. (**A**) Representative pseudo-colored mIHC images showing CD206, F4/80 (macrophages), and CD3 (T cells) in KPC WT tumors (left) and in HuR-KO tumors (right) and (**B**) summary data depicting averaged immune cell density in KPC WT (n=4) and HuR-KO tumors (n = 4). (**C**) ROC curve with corresponding AUC for the machine learning model. (**D**) Model coefficients for the top 18 features that drove the machine learning model to predict tumor genotype WT (black) or HuR-KO (blue). Features are ranked by importance of TME feature type to distinguish tumor genotype. (**E**) Summary data depicting densities of Ki67^+^CD206^-^ macrophages and PD1^+^CD4^+^ T cells and (**F**) summary data depicting densities of Ki67^+^neutrophils and PD-L1^+^ CD11b^-^ macrophages in KPC WT (n= 20 tissue regions) and HuR-KO tumors (n = 11 tissue regions). (**G**) Stacked bar chart showing the average cellular composition of each of eight RCNs. Error bars represent standard error of the mean. *P < 0.05; **P < 0.01; ***P < 0.005; ****P < 0.001, Mann–Whitney U-test to determine statistical significance (**E,F**).

Our machine learning model accurately classified tumor genotype (WT vs HuR-KO) as evidenced by an area under the receiver operating characteristic curve (AUC) of 0.97, with additional performance metrics all scoring above 0.83 (**Fig. 4C** and Supplementary Fig. S2B). This high model performance enabled us to delve into the model coefficients representing the weighted combination of top TME features differentiating WT versus HuR-KO tumors. Of the total 98 TME features, the top 18 features discriminating WT tumors from HuR-KO tumors accounted for 80% of all coefficient weights driving model predictions, and 15 of these top 18 features were statistically significant (**Fig. 4D** and Supplementary Fig. S2C); together, these results indicate these features represent the majority of significant TME differences between WT and HuR-KO tumors.

Consistent with our flow cytometry data (**Fig. 3A**), and despite the relatively low amount of T cells in the total cell population, the model identified increased CD4^+^ T cell density in the HuR-KO tumors as compared to the WT tumors (**Fig. 4B** and **D**). HuR-KO tumors also possessed increased densities of monocytes as compared to the WT tumors. Interestingly, from a spatial perspective, the model detected an increase in cell–cell proximity (defined by 20µm in distance) between CD206^-^ macrophages and CD4^+^ T cells in HuR-KO tumors, while WT tumors had an increase in spatial proximities predominately involving neutrophils, CD206^+^ macrophages and CD11b^-^ macrophages, monocytes, and neoplastic epithelium (**Fig. 4D** and Supplementary Fig. S2D). Delving deeper into these top spatial features highlighted by the model, we found higher densities of Ki-67^+^CD206^-^ macrophages (anti-tumoral “M1-like”) and PD-1^+^CD4^+^ T cells in HuR-KO tumors, which aligns with higher antitumor immunity in HuR-KO tumors (**Fig. 4E**). On the contrary, we found higher densities of Ki-67^+^neutrophils and PD-L1^+^ CD11b^-^ macrophages in WT tumors, which aligns with an immunosuppressive phenotype in WT tumors (**Fig. 4F**).

The model also identified four RCNs as being main differentiators between WT and KO tumors. Briefly, RCNs represent phenotypically distinct spatial groupings—or neighborhoods—of cells within a 60 µm radius of each other that are present across tumor samples. Cells were clustered according to their neighborhood composition and were assigned to one of eight RCNs, each with unique phenotypic compositions (**Fig. 4G**). The machine learning model identified that RCNs 3 and 6 (predominantly neoplastic epithelium and CD11b^-^ macrophages) were enriched in KPC WT tumors, while RCNs 4 and 5 were enriched in HuR-KO tumors (**Fig 4D** and Supplemental Fig. S2E). RCN 4 had the most unique clustering of T cells, CD206^-^ macrophages, and monocytes (**Fig. 4G**), indicating a potential lymphoid-myeloid communication relationship that supports T cell function in an HuR-dependent manner. Together, the mIHC results indicate that the TME phenotypic and spatial landscape differs between HuR-deficient and HuR-competent PDAC tumors. Specifically, our results indicate HuR mediates immune suppression by limiting T cell infiltration and activation, and instead promotes a myeloid-driven immunosuppressive spatial landscape.

### HuR restricts the effector functions of CD4^+^ and CD8^+^ T cells in PDAC

Drawing from our observation that tumor-intrinsic HuR suppressed the activation levels of PDAC-infiltrating T cells, we next asked whether HuR constrained the acquisition of effector functions of tumor-infiltrating T cells. To that end, we performed direct *ex vivo* reactivation of T cells infiltrating WT or HuR-KO PDAC tumors and captured and assessed their cytokine production via intracellular cytokine staining quantified by flow cytometry. By unsupervised clustering all CD4^+^ T cells based on their cytokine production, CD4^+^ T cells derived from WT and HuR-KO tumors naturally separated in two clusters demonstrating their distinct phenotypes (**Fig. 5A**). Consistent with increase in activation markers, CD4^+^ T cells derived from HuR-KO tumors exhibited increase in cytokine production and polyfunctionality (**Fig. 5A** and **B**). Additionally, CD8^+^ T cell derived from HuR-KO tumors displayed an increase in IFN-γ production compared to CD8^+^ T cells derived from WT tumors (Supplementary Fig. S3A).

**Figure 5.**
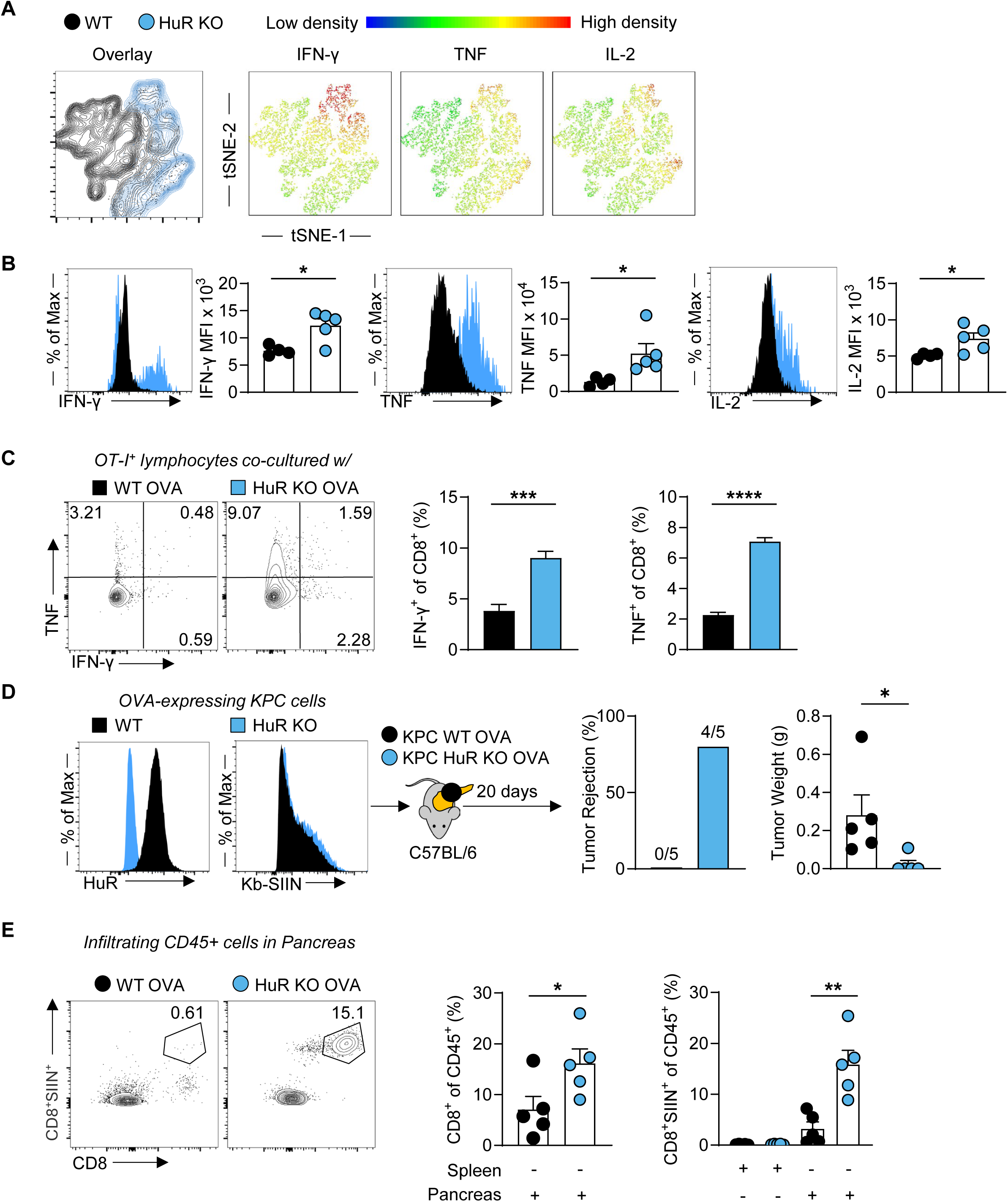
HuR restricts the function and infiltration CD4 and CD8 T cells in PDAC. (**A**) Concatenated single-cell overlay and colorimetric density tSNE based depiction of the indicated proteins by flow cytometry, and (**B**) representative and summary quantification of cytokines following *ex vivo* stimulation of isolated CD4^+^ T cells from KPC WT and HuR-KO tumors (n = 5). (**C**) Representative flow cytometry plot and summary data depicting cytokine productions of OT-I CD8^+^ T cells co-cultured with KPC WT OVA or KPC HuR-KO OVA tumor cells (n = 3). (**D**) Representative flow cytometry plot depicting HuR expression and Kb-SIIN staining of KPC WT OVA or KPC HuR-KO OVA tumor cells prior to orthotopic implantation, and summary data of tumor rejection rate and final tumor weight at day 20. (**E**) Representative flow cytometry plot and summary data enumerating spleen and pancreas infiltrating CD8^+^ T cells and SIIN^+^ CD8^+^ T cells from KPC WT OVA and KPC HuR-KO OVA tumor-bearing mice (n = 5).| Error bars represent stand deviation (**C**) and standard error of the mean (**B,D,E**). *P < 0.05; **P < 0.01; ***P < 0.005; ****P < 0.001, unpaired 2-tailed Student’s t tests.

### HuR restricts the infiltration of antigen-specific CD4^+^ and CD8^+^ T cells in PDAC

To determine the impact of HuR loss in PDAC on the function and abundance of tumor-specific T cells, we retrovirally transduced KPC cells with an ovalbumin (OVA)-expressing construct to make KPC cells express full-length OVA for processing and presentation on MHC (**Fig. 5D**, left). When *ex vivo* expanded and activated ovalbumin-specific CD8^+^ T cells (OT-I cells) were co-cultured with KPC OVA cells, we found that KPC WT OVA cells exhibited suppression of OT-1 T cell cytokine production (IFN-γ and TNF-α production) relative to KPC HuR-KO OVA cells (**Fig. 5C**), despite there being no difference in MHC-I abundance or antigen presentation in KPC HuR-KO cells in response to interferon stimulation (**Fig. 5D**). This co-culture model continues to support the notion that tumor-intrinsic HuR suppresses local T cell function in a manner independent of antigen presentation or interferon response. To evaluate the endogenous T cell responses against the OVA antigen in the presence or absence of tumor-intrinsic HuR, HuR WT or HuR-KO OVA cells were orthotopically implanted into C57BL/6 mice. Notably, KPC HuR-KO OVA tumors were rejected in 80% of mice, with significantly smaller tumors than KPC WT OVA tumors in those instances where a tumor was indeed evident (**Fig. 5D**). Given the absence of a pancreatic tumor in most instances of HuR-KO KPC OVA, we evaluated the peritumoral tissues in both groups of tumor-bearing hosts. We found statistically significant increases in the amount of CD8^+^ T cells, and OVA tetramer^+^ CD8^+^ T cells in hosts bearing HuR-KO OVA tumors than WT OVA tumors (**Fig. 5E**). We observed much lower SIINFEKL tetramer staining in the spleen of tumor bearing mice, and no difference in the amounts of OVA tetramer^+^ CD8^+^ T cells between spleens from WT tumor-bearing hosts compared to HuR-KO tumor-bearing hosts (**Fig. 5E**). Together these data indicate that tumor-intrinsic HuR suppresses the number of functional populations (cytokine producing, and antigen-specific) of T cells in the PDAC TME, thus promoting tumor immune evasion.

### HuR stabilizes anabolic transcripts and metabolic pathways

To identify the mechanism underlying HuR-dependent immune suppression and tumor progression in our immunocompetent model, we performed bulk-RNA sequencing on KPC WT and HuR-KO cells (Supplementary Fig. S4A). GSEA analysis identified established signatures involved in MYC pathways and metabolic reprogramming, including adipogenesis and cholesterol homeostasis, were higher in the KPC WT cells compared to the HuR-KO cells (Supplementary Fig. S4A and B). Given that HuR is known to stabilize RNA by directly binding to the 3’UTR regions of transcripts, we sought to identify those RNAs in the KPC WT cells using ribonucleoprotein immunoprecipitation (RIP) followed by high-throughput RNA sequencing (**Fig. 6A**). We validated HuR-RIP pulldown with immunoblot analysis from the bead-bound lysate and flow-through. Validating our protocol, motif analysis showed preferential binding to AU-rich sequences as previously described for HuR (**Fig. 6C** and Supplementary Fig. S4C)^47^. Additionally, established HuR metabolism targets, the *IDH1* 3’ UTR^13^, and the *Rictor* 3’ UTR, demonstrated significantly peak enrichment in comparison to reference control *GAPDH* loci (**Fig. 6B** and Supplementary Fig. S4D). KEGG pathway analysis of the 1847 bound mRNAs revealed enrichment of transcripts involved in metabolism, including MAPK signaling, mTOR signaling, HIF1-α pathway and proteoglycan pathways, consistent with a model wherein HuR drives PDAC nutrient consumption to the potential detriment of adjacent cell populations (**Fig. 6D**).

**Figure 6.**
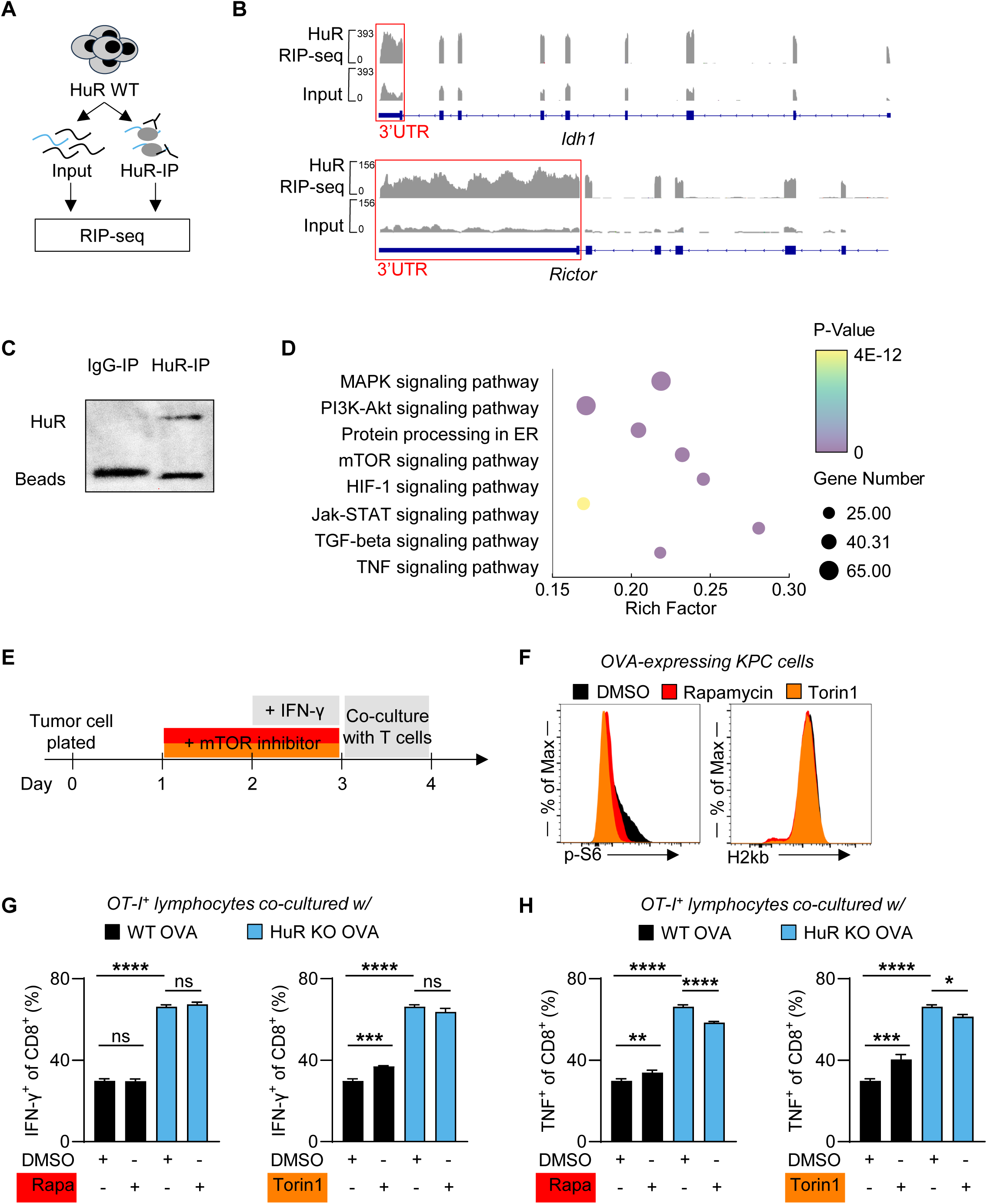
HuR stabilizes anabolic transcripts and metabolic pathways. (**A**) Schematic depicting RIP-seq process to identify HuR-bound transcript (n = 3). (**B**) RIP-seq reads genome alignment plot to *IDH1 and Rictor* loci (3’UTR region highlighted with red box). (**C**) Representative immunoblot validation of HuR pulldown in KPC cells in comparison to IgG control. (**D**) Summary depiction of enrichment scores (RichFactor) of KEGG pathways with transcripts comparatively enriched in HuR-IP samples. (**E**) Schematic depicting the pre-treatment of tumor cells with mTOR inhibitors (Rapamycin and Torin1) and IFN-γ before co-culture with OT-I T cells. (**F**) Representative flow cytometry plot depicting *p*-S6 and H2-Kb staining of KPC WT OVA cells after mTOR inhibitors and IFN-γ treatments. (**G**, **H**) Summary data depicting cytokine productions of OT-I CD8^+^ T cells co-cultured with pre-treated KPC WT OVA or KPC HuR-KO OVA tumor cells (n = 3). Error bars represent stand deviation (**G**, **H**). *P < 0.05; ***P < 0.005; ****P < 0.001, one-way ANOVA test.

To test our hypothesis that HuR increases mTOR activity and drives metabolic competition to suppress local T-cell function, we pretreated KPC OVA cells with mTOR inhibitors (AZD8055 and Torin1) and IFN-γ, and then co-cultured the pretreated KPC OVA cells with OVA-antigen specific OT-I cells (**Fig. 6E**). The pretreated KPC OVA cells had decreased mTOR activity by the measurement of decreased levels of phosphorylated-ribosome subunit 6 (p-S6). Of note, we validated that the inhibitors did not impair H2kb levels in these tumor cells (**Fig. 6F**). Confirming our hypothesis, we observed that OT-I T cells co-cultured with mTOR-inhibited KPC WT OVA cells had increased cytokine production (IFN-γ and TNF) in comparison to T cells co-cultured with vehicle-treated KPC WT OVA cells (**Fig. 6G** and **H**). In comparison, T cells co-cultured with KPC HuR-KO OVA cells demonstrated higher cytokine production *de novo,* independent of additional mTOR inhibition – suggesting overlap between the two pathways (**Fig. 6G** and **H**). This data further supports our hypothesis that HuR-KO tumor cells have lower mTOR activity at baseline and as a result, less nutrient competition with T cells, thus mTOR inhibition in HuR-KO tumor cells could not increase T cell function further. In addition, because of the different inhibition capacity of different inhibitors, T cells co-cultured with tumor cells treated with mTOR1 and mTOR2 inhibitor, Torin1, displayed increase in both IFN-γ and TNF production, while mTOR1 inhibitor, rapamycin, only displayed increase in TNF production (**Fig. 6G** and **H**). Taken together, these data indicate mechanistically tumor-intrinsic HuR directly binds to and stabilizes transcripts involved in mTOR pathways, and the HuR-mediated increase of mTOR activity in tumor metabolic competes with local T cells and ultimately suppress T cell function.

### PDAC resistance to Immune checkpoint blockade is dependent on HuR

As previous discussed, ICB showed limited response in PDAC patients with deficiency in DNA mismatch repair machinery^7^, which suggested that the PDAC TME is inherently immune suppressive. Drawing from the observation that PDAC HuR is responsible for constraining the infiltration and function of tumor specific T cells, we next asked whether PDAC therapeutic resistance to ICB treatment was dependent on HuR function. Following a published treatment regimen^3^, we treated KPC WT and HuR-KO tumors-bearing mice with αPD-1 or control αIgG (**Fig. 7A**). Consistent with prior reports, αPD-1 treatment alone in KPC WT tumors did not lead to tumor regression. However, αPD-1 treatment in KPC HuR-KO tumors induced significant and almost total tumor regression (**Fig. 7A**). The αPD-1 treatment did not induce significant loss of body weight in mice, indicating the treatment was well-tolerated in both groups (**Fig. 7B**). Consistent with previous observations (**Fig. 3A**), HuR depletion led to increased CD4^+^ T cell infiltration in tumors. While αPD-1 treatment in HuR-KO tumors did not further boost the CD4^+^ T cell infiltration compared to the IgG treatment, interestingly, αPD-1 treatment in HuR-KO tumors induced higher infiltration of CD8^+^ T cells with no effect observed in the WT tumors (**Fig. 7C**). Together, these data indicate that CD8^+^ T cells are the main responder to αPD-1 treatment in HuR-KO tumors, and support our hypothesis that HuR depletion in PDAC tumors provides a greater proportion of T cells in the TME, which increased the capacity for enhanced antitumor effector activity.

**Figure 7.**
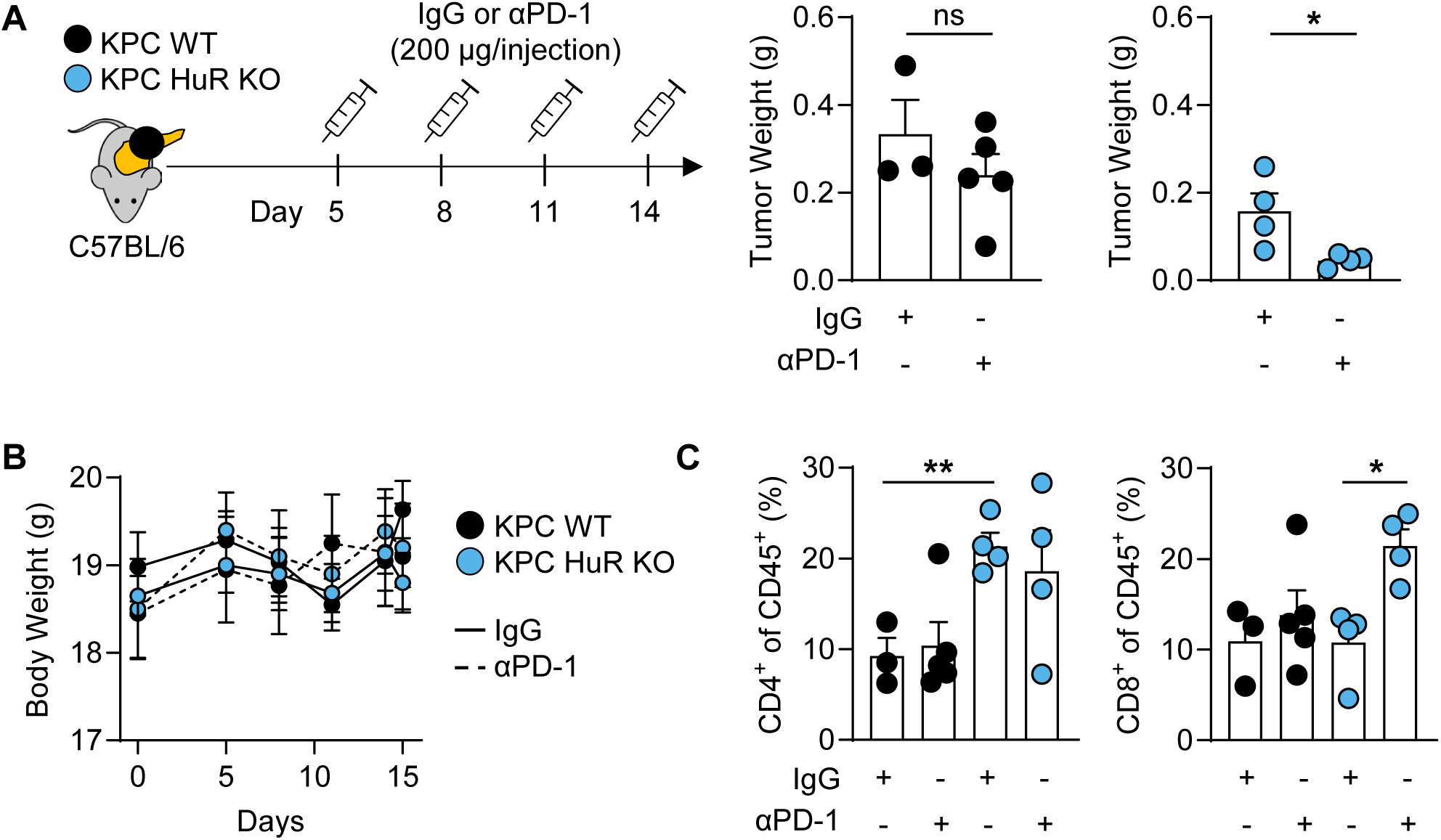
PDAC resistance to immune checkpoint blockade is dependent on HuR. (**A**) Schematic for αPD-1 treatment on KPC tumor bearing mice, and summary data on final tumor weight at day15. (**B**) Body weight of mice in (**A**) by group tracked over time (n = 4). (**C**) Summary analysis enumerating tumor infiltrating lymphocytes (TIL) of KPC WT and HuR-KO tumors in mice received IgG or αPD-1 treatment in (**A**). Error bars represent standard error of the mean. *P < 0.05; **P < 0.01, unpaired 2-tailed Student’s t tests (**A**), one-way ANOVA test (**C**).

## DISCUSSION

The immune landscape of PDAC TME is highly immunosuppressive and is a major barrier to effective immunotherapy response^2,8^. Hence, understanding the underlying molecular mechanisms is key to advancing immunotherapy strategies. From multi-omics analysis on large patient databases, the RNA-binding protein, HuR, negatively correlates with both patient survival and total lymphocytes infiltration in PDAC. Herein, we found that HuR expressed by PDAC promotes tumor growth in the immunocompetent PDAC model. Within PDAC cells, HuR directly binds to and stabilizes anabolic transcripts, ultimately limiting the abundance and function of CD4^+^ and CD8^+^ T cells. As the primary mediators of antitumor immunity, the T cells in the HuR-proficient PDAC TME had limited effector functions in cytokine production and antigen-specific response. This mechanism could be exploited with therapeutic potential, as depletion of HuR sensitizes PDAC tumor to ICB treatment with increased efficacy.

In previous studies, loss of HuR in human PDAC xenografts in immunocompromised hosts had no impact on tumor growth^24^, which complements our findings that HuR has an immune-mediated tumor suppressive role. In addition, previous work targeting overexpressing HuR in the mouse pancreas led to chronic inflammation in the TME and promoted tumor development, indicating HuR contributes to an unbalanced inflamed environment^23^. Herein, this is the first report of PDAC tumor-intrinsic HuR having a potent extrinsic effect on the immune cells in the TME. This new discovery of HuR function fits with its known role in supporting tumor initial and tumor progression, and future study should investigate how HuR-mediated immune suppression promotes PDAC initiation from pancreatitis. Similarly, another key transcriptional regulator in PDAC, Myc, has been shown to control immune evasion in PDAC^11^. Therefore, studying pro-oncogenic molecules like HuR and Myc in immune-competent models may provide a much-needed understanding of the post-transcriptional and transcriptional tumor-intrinsic networks involved in immune suppression in the PDAC TME.

Interestingly, we did not observe a high correlation between gain/loss of RNA transcripts by RNA-sequencing and alignment abundance on RNA-immunoprecipitation. Overlapping these HuR-bound targets with those whose expression was reduced upon HuR-KO in the same cell lines identified 141 RNAs that are dependent on HuR binding to maintain transcript levels (data not shown). While the low percentage of overlap was surprising, published works have observed similar non-linear relationships between RNA-binding proteins and their targets^47,48^. These results further illustrate the complicated regulation mechanisms of RNA-binding proteins with its binding targets in general. A more granular understanding of the relationship between HuR 3’ UTR binding and the impact on protein abundance will inform our ability to leverage HuR function for translational purposes.

Transcriptomic analyses in our KPC model confirmed multiple dysregulated pathways, which have been previously illustrated to be regulated by HuR^16,17^. Specifically, HuR directly upregulates multiple pathways in tumor remodeled metabolism^13^. Metabolic reprogramming in the tumor through the Warburg effect, characterized by aerobic glycolysis, not only enhances tumor proliferation but also depletes the limited resources within the TME^14,49^. Recent metabolite profiling of the PDAC interstitial fluid showed altered nutrient availability and nutrient deprivation in the PDAC TME^50^, suggesting metabolic competition could be one of the mechanisms that tumor cells use to suppress immune cell function. Particularly, reports have shown that activated T cells need to reprogram their glucose and lipid metabolism to support their effector functions^51,52^, and Tregs are more adapted to glucose deprivation with an increase in fatty-acid oxidation^53^. These required nutrients are in concordance with the HuR-mediated dysregulated metabolic pathways, suggesting HuR mediates immune suppression via metabolic competition in the PDAC TME. In lieu of this, previous finding that PDAC WT and HuR-KO tumors grow similarly in immunocompromised model further supports the notion that HuR-mediated anabolic pathways do not support tumor growth by proliferation^24^, instead by inducing immune suppression. In total, our results warrant further exploration to define the mechanistic connection between HuR-mediated PDAC metabolism and T cell function and differentiation in the TME.

In the PDAC immune landscape, our findings once again demonstrate the potential of CD8^+^ T cells to be harnessed for PDAC tumor control, while also highlighting that CD4^+^ T cells can play a crucial role in addition to well-described CD8^+^ T cells. Similar observations of CD4^+^ T cells having anti-PDAC response were found in the setting of αCD40 stimulation^54^. These findings support the potential of targeting CD4^+^ T cells to augment anti-tumor responses. In addition to impacting T cells, our mIHC analysis demonstrated that tumor-intrinsic HuR may have a broader impact on the immune landscape of the PDAC TME, impacting the spatial organization and proliferation status of multiple myeloid lineages, including increased proliferation of neutrophils and decreased proliferation of anti-tumoral “M1-like” macrophages. PDAC-infiltrating neutrophils have been implicated to associate with poor prognosis in patients^55,56^, and our findings reveal a potential novel upstream tumor-mediator regulator of neutrophil proliferation in PDAC. Still, the implications of these findings highlight the need for further investigation into the mechanism of HuR mediated myeloid cell activities.

Broadly, these findings advance the paradigm that tumor-intrinsic mechanisms limit T cell infiltration and the efficacy of anti-tumor immunity within the TME. Specifically, we highlight a novel role of tumor-intrinsic HuR as a modulator of PDAC immune evasion. Currently, targeting strategies against HuR are under development (e.g., siHuR nanotherapies and small molecule inhibitors)^57^, and could potentially break PDAC resistance to immune-based anti-cancer therapies (i.e., checkpoint blockade and T cell transfer) as demonstrated here in our pre-clinical KPC model. Because HuR also has cell-intrinsic functions in intratumoral immune populations, further work will be required to understand the combined impact of HuR inhibition on tumor control.

## ACKNOWLEDGEMENTS

We would like to thank all members of the Brody laboratory and the Eil laboratory, the Brenden-Colson Center for Pancreatic Care, and the Knight Cancer Institute for their continued support and discussions regarding this work. This work was supported by the National Cancer Institute (NCI) of the National Institutes of Health (NIH) under awards R01 CA212600 (to J.R.B), U01 CA224012 (to R.C.S. & J.R.B), R21 CA263996 (to R.C.S. & J.R.B), and R01GM147365 (to Z.X.), K08 CA256179 (to R.E.), and P30 CA069533 (to L.M.C.), American Association for Cancer Research (AACR) under award 15-90-25-BROD (to J.R.B) and Transformative Oncology Research (to R.E.), the Hirshberg Foundation, and the Brenden-Colson Center for Pancreatic Care, American Society of Clinical Oncology (ASCO) Career Development Award (to R.E.), PanCAN Career Development Award (to R.E.), America Cancer Society, the Susan G. Komen Foundation, the National Foundation for Cancer Research, and Hildegard Lamfrom Endowed Chair in Basic Research (to L.M.C.). We thank and acknowledge the invaluable work performed by members of the OHSU Research Cores and Shared Resources including members of the Flow Cytometry Core, Advanced Light Microscopy Core, and Histopathology Shared Resource. We thank the OHSU Department of Comparative Medicine for their support in animal care, especially Allie Buckner for her hands on care of our animals.

## AUTHOR CONTRIBUTIONS

Conceptualization: Y.G., J.R.B., R.E.; resource: A.G.; software: K.E.B., K.H., C.C.; formal analysis: Y.G., K.E.B., S.S., K.H., C.C.; supervision: A.G., Z.X., L.M.C., R.C.S., J.R.B., R.E.; funding acquisition: A.G., Z.X., L.M.C., R.C.S., J.R.B., R.E.; investigation: Y.G., J.M.F., A.Q.B., S.S., N.K., N.B., G.A.M.; visualization: Y.G., S.S.; writing – original draft: Y.G., J.R.B., R.E.; writing – review & editing: Y.G., J.M.F., A.Q.B., S.S., K.E.B., L.M.C., R.C.S., J.R.B., R.E..

---

**Supplement Figure 1.**
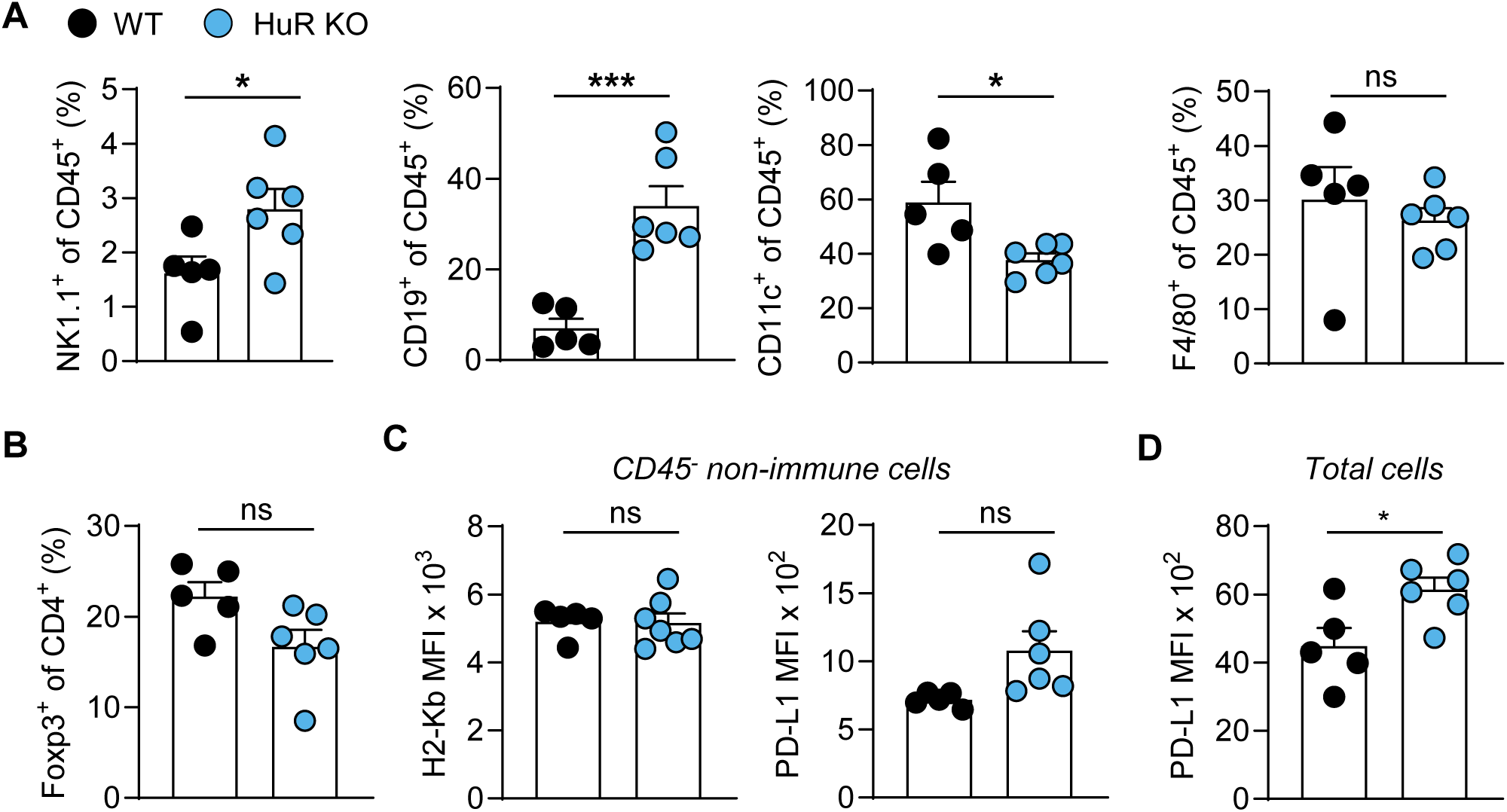
Tumor-intrinsic HuR mediates immune composition in the PDAC TME. (**A**) Summary analysis enumerating tumor infiltrating NK cells, B cells, dendritic cells, macrophages, and (**B**) Foxp3^+^ CD4 T-cell percentages. (**C**) Summary data depicting H2-kb and PD-L1 expression on non-immune cells, and (**D**) PD-L1 expression on all cells in orthotopic KPC WT and HuR-KO tumors (n = 5). Error bars represent standard error of the mean. *P < 0.05; ***P < 0.005, unpaired 2-tailed Student’s t tests.

**Supplement Figure 2.**
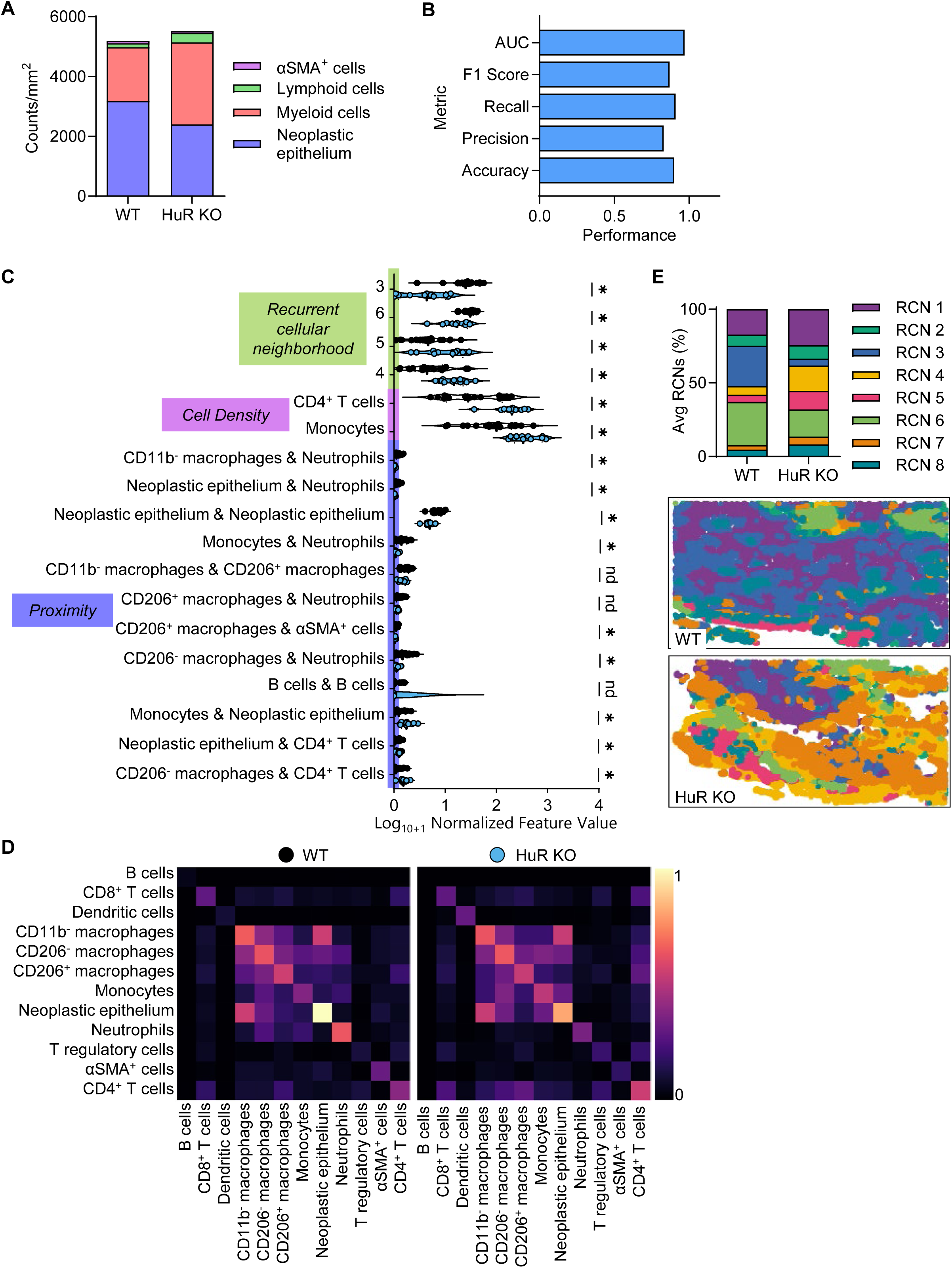
HuR limits T cell infiltration and activation, promoting an immunosuppressive and myeloid-driven TME landscape. (**A**) Summary data depicting average stromal, immune, and neoplastic epithelium cell density in KPC WT (n=4) and HuR-KO tumors (n = 4). (**B**) Bar chart showing performance metrics for the machine learning model. (**C**) Violin plot showing the normalized values of the top 18 features driving the model. Features are organized by type on y-axis and log_10+1_ normalized values are shown on the x-axis. (**D**) Heatmaps showing average normalized frequency of spatial proximities between two cell types for each tumor group. (**E**) Summary data showing average fraction of RCNs present in KPC WT (n=4) and HuR-KO (n=4) tumors and scatterplot reconstructions of representative tissue regions with each cell colored by its RCN designation. Error bars represent standard error of the mean. *P < 0.05, Mann–Whitney U-test to determine statistical significance (**C**).

**Supplement Figure 3.**
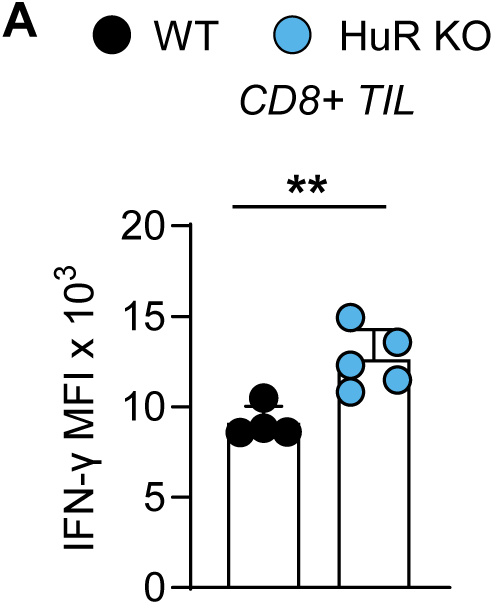
HuR restricts the function and infiltration CD4 and CD8 T cells in PDAC. (**A**) Summary quantification of IFN-γ production following *ex vivo* stimulation of isolated CD8+ T cells from KPC WT and HuR-KO tumors (n = 5). Error bars represent standard error of the mean. **P < 0.01, unpaired 2-tailed Student’s t tests.

**Supplement Figure 4.**
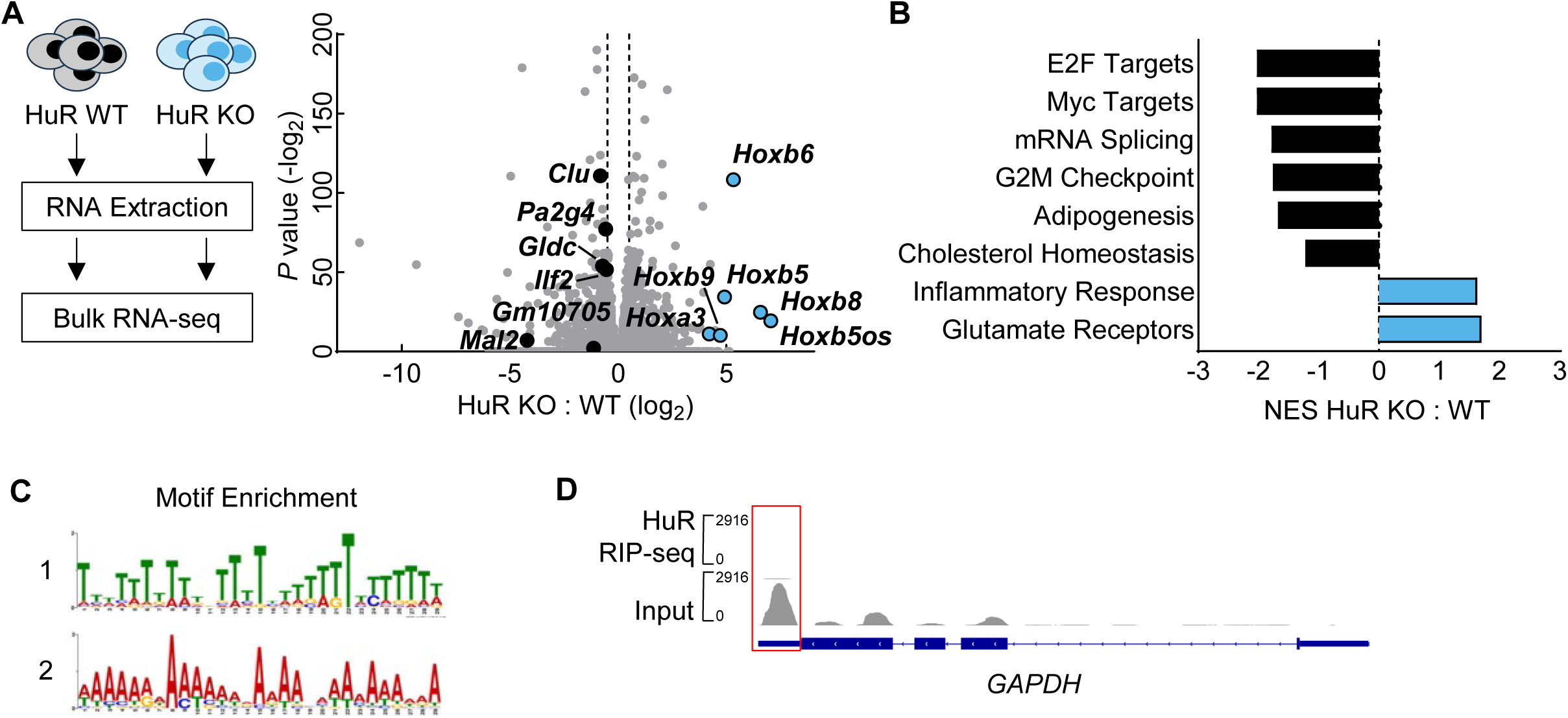
HuR stabilizes anabolic transcripts and metabolic pathways. (**A**) Schematic and volcano plot depicting whole-transcriptomic changes between KPC WT and KPC HuR-KO tumor cells (n = 3). (**B**) Summary depiction of normalized enrichment scores (NES) of gene-sets with transcripts comparatively enriched or lost between indicated groups. (**C**) Top 2 motif enrichment sequences from HuR-IP targets. (**D**) RIP-seq reads genome alignment plot to *GAPDH* locus (3’UTR region highlighted with red box). FDR q-value < 0.05, weighted Kolmogorov–Smirnov statistic test (**B**).

**Supplement Table 1.** Antibody information for flow cytometry staining.

| **Marker** | **Clone** | **Fluorescence** |
| --- | --- | --- |
| CD11c | N418 | BV650 |
| CD19 | 1D3 | BV711 |
| CD25 | PC61 | BV421 |
| CD4 | RM4-5 | PCP-Cy5.5 |
| CD4 | RM4-5 | BUV805 |
| CD45 | 30-F11 | AF700 |
| CD62L | MEL-14 | BUR395 |
| CD69 | H1.2F3 | APC-Cy7 |
| CD8 | 53-6.7 | FITC |
| CD8 | 53-6.7 | APC-Cy7 |
| CD90.2 | 53-2.1 | BV786 |
| F4/80 | BM8 | BV421 |
| FasL | MFL3 | PE |
| Foxp3 | FJK-16s | PE |
| GranzB | NGZB | PE |
| IFNg | XMG1.2 | APC |
| IL-2 | JES6-5H4 | FITC |
| Ki67 | SolA15 | BUV805 |
| H2-kb | AF6-88.5 | FITC |
| I-A,E | M5/114.15.2 | APC-Cy7 |
| NK1.1 | PK136 | PE-Cy7 |
| PD-1 | RPM1-30 | APC |
| PD-L1 | MIH5 | APC |
| TBET | 4B10 | PE-Cy7 |
| Tim3 | 5D12 | BV650 |
| TNF | MP6-XT22 | BV421 |
| LiveDead |  | Aqua |

**Supplement Table 2.**
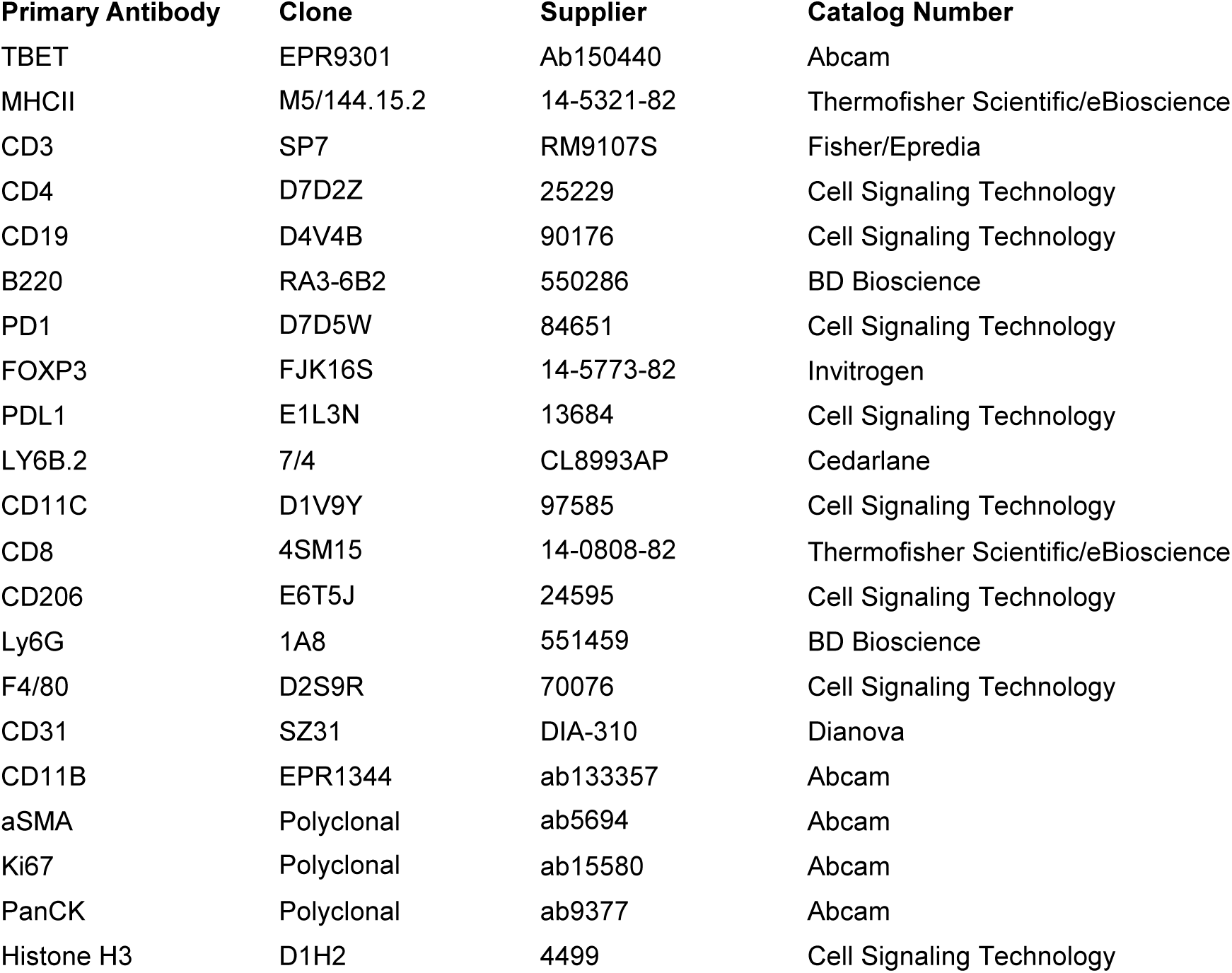
Antibody information for mIHC staining.

**Supplement Table 3.**
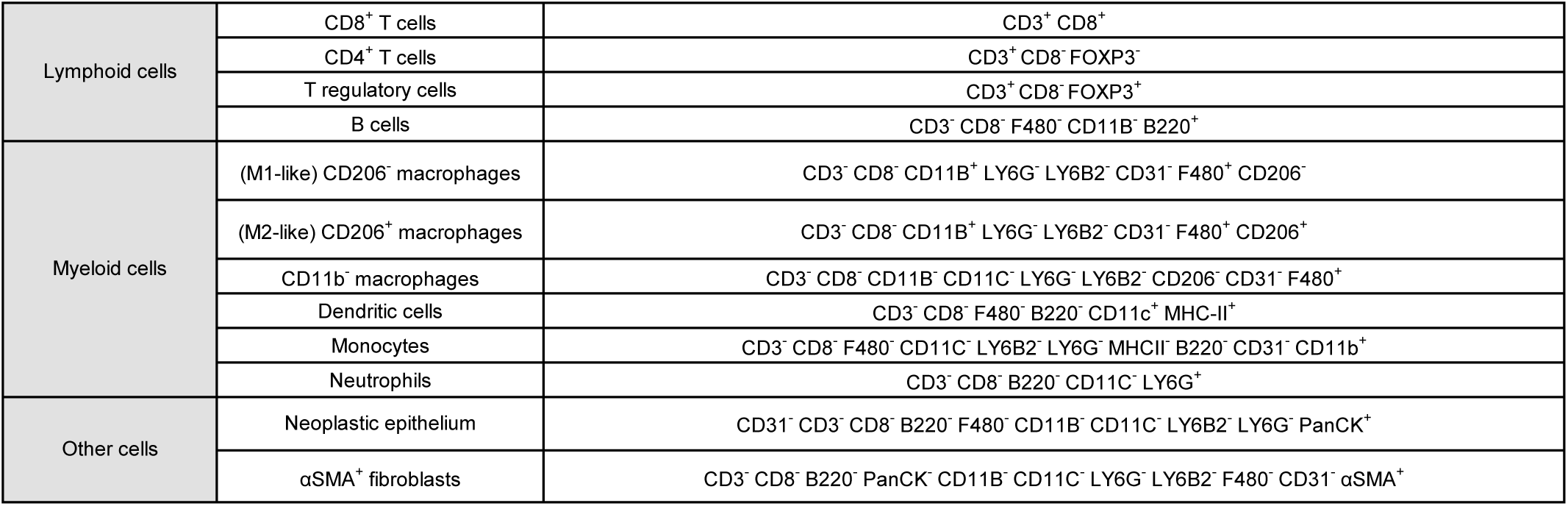
Gating strategies for lineage assignment in mIHC.

